# Emergence of histone-based chromatin complexity in Asgard archaea

**DOI:** 10.64898/2026.07.02.735847

**Authors:** Chung Hyun Cho, Colleen S. Russett, Zachary H. Harvey, Nika Pende, Logan H. Hodgskiss, Jérémy Cangi, Philipp Radler, Karin Hager, Miroslav Homola, Richard Imre, Gabriela Krssakova, Shun’ichi Ishii, Harsh M. Ranawat, Isabelle Anna Zink, Nevena Maslać, Joshua N. Hamm, Anja Spang, Masaru K. Nobu, Silvia Bulgheresi, Svetlana O. Dodonova, Hiroyuki Imachi, Elisabeth Roitinger, Nicholas A. T. Irwin, Florian K. M. Schur, Christa Schleper, Frédéric Berger

## Abstract

The emergence of the eukaryotes coincided with the diversification of histone proteins and their post-translational modifications by enzymes that constitute the core of eukaryotic chromatin. Yet the evolutionary origins of this regulatory machinery are unknown. Here, we show that the key molecular components of histone-based chromatin regulation are present in the Asgard archaea, the closest prokaryotic relatives of eukaryotes. Asgard histones are abundant and have extended N-terminal tails rich in lysine residues that can be post-translationally modified, all of which are features shared with eukaryotic histones. In line with these findings, we identify enzymes from Asgard archaea that deposit or remove lysine acetylation on histone tails in vitro. Moreover, Asgard sirtuin deacetylases (SIR2 proteins) restore chromatin silencing in yeast, demonstrating the functional compatibility of Asgard enzymes with eukaryotic histone substrates. Our findings establish that the foundations of histone-based chromatin predate eukaryogenesis and place Asgard archaea as an evolutionary intermediate in the emergence of eukaryotic chromatin.

## Introduction

Histone-based chromatin regulation is a hallmark of eukaryotic complexity. The association of histone proteins with DNA condenses the genome and serves as a regulatory scaffold for gene expression, genome replication and maintenance, and cellular differentiation^1,2^. The fundamental element of eukaryotic chromatin is the nucleosome, comprising DNA wrapped around an octamer of histones H2A, H2B, H3, and H4, whose disordered tails are subject to a diverse array of histone post-translational modifications (hPTMs) deposited and removed by histone-modifying enzymes^3,4^. These modifications, including the acetylation and methylation of specific lysine residues, constitute a ‘histone code’ that regulates transcription and genome organization^5–9^. Histone-like proteins have been identified in various prokaryotes and viruses, and histones in Archaea are known to be involved in transcriptional regulation and environmental adaptation^10–16^. Although prokaryotic histones bind and compact DNA, site-specific hPTMs have not been described outside of eukaryotes, and the evolutionary origins of histone-based genome regulatory machinery in eukaryotes are unclear^17–21^.

The discovery of Asgard archaea (also formally known as kingdom *Promethearchaeati*^22^ or phylum *Asgardarchaeota*^23^) reshaped our understanding of the tree of life, firmly placing the origin of eukaryotes within the archaeal domain^24–26^. Furthermore, the recent cultivation of Asgard archaea has revealed that they harbor certain cellular features resembling eukaryotic complexity, including cell protrusions formed by cytoskeletal proteins homologous to eukaryotic actin and tubulin^27–30^. This evidence is consistent with genomics studies that have revealed that Asgard archaea possess an extensive repertoire of eukaryotic signature proteins^24^, including genes that encode proteins with features of eukaryotic chromatin^19,31^. Asgard histones have been shown to form hypernucleosomal chromatin assemblies in vitro^32^, however, it remains unknown whether Asgard archaea in vivo possess histone-based chromatin or the regulatory machinery required to write and erase hPTMs.

Whether chromatin capable of regulation through hPTMs originated within the eukaryotic lineage or has deeper prokaryotic roots has remained an open question. To address this point, we obtain a large proteomics dataset representative of archaeal diversity and biochemical analyses to characterize histones and histone-modifying enzymes in Asgard archaea. We show that Asgard archaea possess abundant and diverse histones with tails and undergo site-specific hPTMs. We demonstrate that enzymes write and erase these modifications in vitro and use functional complementation assays in yeast. Altogether our findings reveal that histone-based chromatin complexity arose long before the origin of eukaryotic cells.

## Results

### Asgard archaea have abundant and diverse tailed histones

Eukaryotic chromatin is defined by histones, which act as the dominant genome-packaging system. For example, histones comprise ∼2–4% of total protein abundance in the proteomes of human cell lines^33^. To determine whether any archaeal lineage has a similar abundance of histones to eukaryotes, we generated quantitative whole-cell proteomes of 29 species and combined these with 26 publicly available proteomes to represent all three domains of life, comprising in total 6 eukaryotes, 3 Bacteria, and 43 Archaea. Specifically, our datasets have representatives of the 6 Asgard archaea, 12 TACK (also known as kingdom *Thermoproteati*), 22 Euryarchaeota (also known as *Methanobacteriati*), and 3 DPANN (also known as kingdom *Nanobdellati*) (Fig. 1a; Supplementary Fig. 1), enabling a comprehensive survey of archaeal features. Ranking chromatin proteins by their protein abundance confirmed that histones are among the most abundant proteins in eukaryotes (Supplementary Table 1).

**Figure 1.**
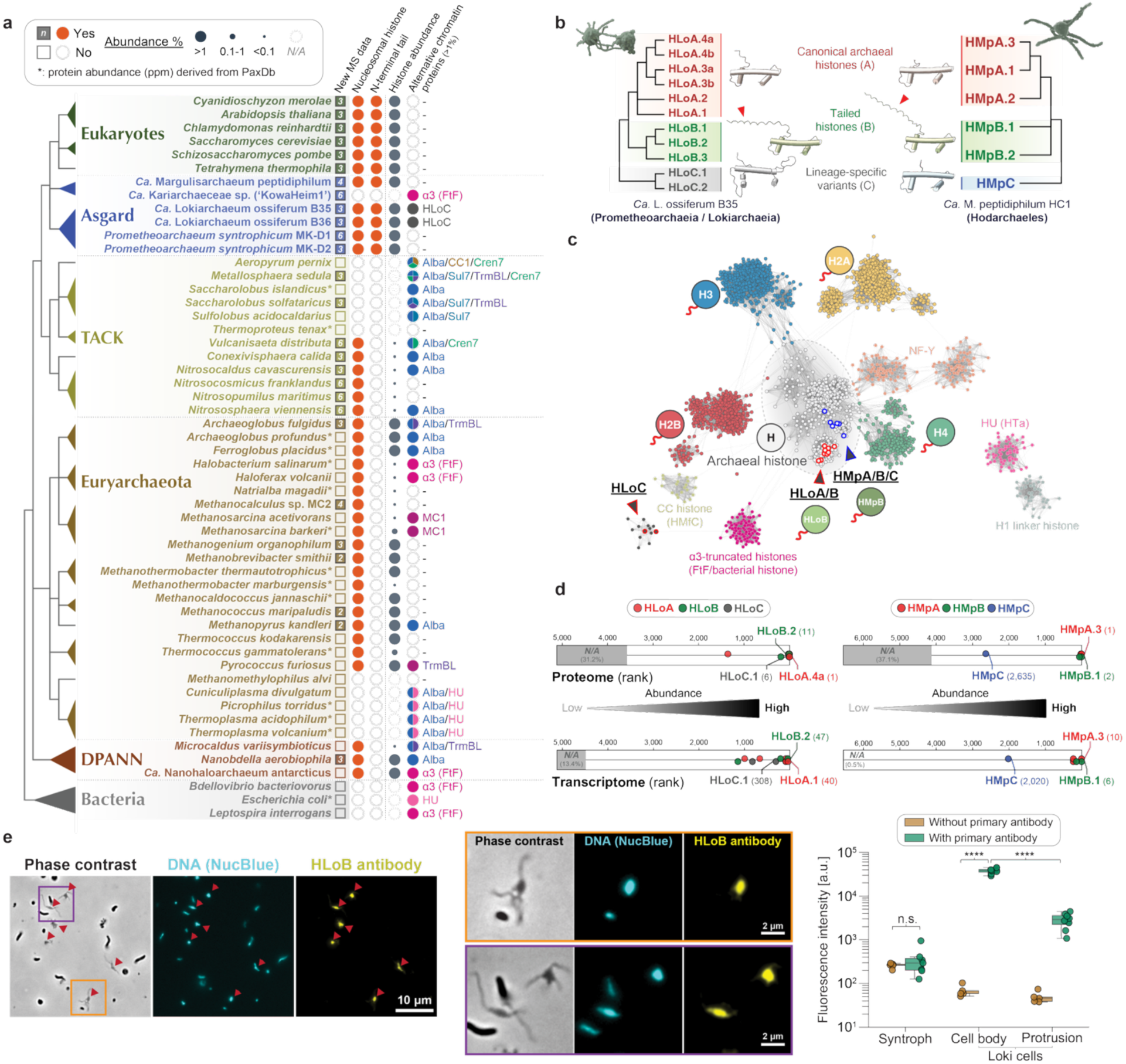
A tree-of-life chromatin analysis reveals eukaryotic-level histone abundance and diversification in Asgard archaea. **a**, Tree-of-life survey of chromatin protein abundance. Whole-cell proteomics of 54 species across bacteria, DPANN, Euryarchaeota, TACK, Asgard, and eukaryotes show the relative abundance of nucleosomal histones and alternative chromatin proteins as a fraction of the total proteome. Proteomics data generated in this study are marked with filled squares (27 species), publicly available datasets are marked with open squares (27 species), and integrated protein abundance from PaxDb is marked with asterisks (15 species). The number of independent mass spectrometry runs (n) is indicated next to each taxon name for filled-square datasets. For alternative chromatin proteins, a threshold of ∼1% total protein abundance was applied based on the eukaryotic chromatinization benchmark. Phylogenetic relationships and nomenclatures were based on an up-to-date archaeal tree of life^62^. iBAQ values are shown in Supplementary Table 1. **b**, Histone diversity in representative Asgard lineages. Simplified phylogeny showing histone families in *Ca.* Lokiarchaeum ossiferum B35 (HLo) and *Ca.* Margulisarchaeum peptidiphilum HC1 (HMp). Histone families include canonical archaeal histones that lack N-terminal tails (HLoA/HMpA, red), N-terminal tailed histones (HLoB/HMpB, green), and other lineage-specific variants (HLoC, grey; HMpC, blue). **c**, Sequence similarity network (SSN) of prokaryotic and eukaryotic histone-fold proteins with HU and histone H1 linker. Nodes are colored by classification. Loki-B35 histones (HLoA/B, HLoC) are denoted by arrows with red outlines and Hod-HC1 histones (HMpA/B/C) by arrows with blue outlines with matching-colored outlines on the corresponding nodes. HLoB and HMpB reside within the archaeal nucleosomal cluster but carry eukaryotic-like features. HLoC occupies a peripheral position. **d,** Tailed histones rank among the most abundant proteins in Asgard. Transcriptome and proteome data from *Ca.* L. ossiferum B35 and *Ca.* M. peptidiphilum HC1 show expression rank and protein abundance, respectively. Tailed histones (HLoB, HMpB) are highly expressed and abundant, ranking among major cellular proteins alongside canonical histones. **e,** Representative overview (left) and zoomed (middle) micrographs of immunostaining experiments on Loki-B35 enrichment culture. The DNA signal stained by NucBlue (cyan) co-localizes with HLoB tailed histones, targeted with an antibody (yellow). The quantification of fluorescence intensities (right) shows no change in the measured intensity of syntrophic organisms ± antibody (p-value = 0.22). Raw fluorescence intensity values (a.u.) are plotted on a log_10_-scaled y-axis. Signal intensity in Loki cells is markedly increased with the primary antibody, demonstrating antibody specificity for HLoB histones (p-value = 4.26 × 10^-9^). The fluorescence intensity signal for tailed histones is highly enriched in the cell body compared to protrusions within Loki cells (p-value = 4.51 × 10^-11^). 5 cells for each condition were quantified in experiments without an antibody and 7 cells in experiments with an antibody. The experiments were performed on two B35 cultures.

We found that histones are found broadly and in high abundance among many archaeal groups, suggesting that they are a typical feature of Archaea (Fig. 1a). To establish a benchmark for genome-packaging competence, we analyzed integrated protein abundance data from *Saccharomyces cerevisiae* spanning multiple orthogonal methods (mass spectrometry, immunoblotting of TAP-tagged proteins, and quantification of green fluorescent protein-tagged proteins)^34^, which indicates that histones comprise 1.3% of all proteins in yeast based on their abundance. Our whole-cell proteomics of eukaryotes—across diverse eukaryotic lineages—showed that histone abundance varies from 1.0% to 24.1% (Extended Data Fig. 1). We thus used this lower bound (∼1% of total protein) as a conservative benchmark for a histone abundance that is sufficient to package genomes. Similar levels of histone abundance are observed in the euryarchaeal *Thermococcus kodakarensis*^35^, which forms hypernucleosomes, thus validating the benchmark established from eukaryotes. In other species of Archaea, e.g. euryarchaeal *Haloferax volcanii*, histones comprise less than 0.1% of total protein (Fig. 1a), consistent with their known roles as transcriptional regulators^11,13^. By contrast, in Asgard archaea with tailed histones, histone abundance exceeded the ∼1% benchmark. In four different strains of cultivated *Promethearchaeia* (hereafter called ‘Loki’; also known as class *Ca.* Lokiarchaeia), histone abundance ranged from 3.8% to over 20.8% and in Hodarchaeales (hereafter called ‘Hod’) *Ca.* Margulisarchaeum peptidiphilum HC1 histones comprised as much as 21.2% of total protein (Extended Data Figs. 2a,b). This phenomenon is not a ubiquitous feature of Asgard archaea, however, because ‘KowaHeim1’ (Supplementary Fig. 2), a member of class *Ca.* Heimdallarchaeia, lacks canonical nucleosomal histones and instead has abundant non-nucleosomal histones, which are α3-truncated histones (i.e., face-to-face histones)^36^ (Fig. 1a).

Correspondingly, we observed a mutual exclusivity in Archaea between the abundance of histones and that of alternative chromatin proteins (Extended Data Fig. 2c, 3). These alternative chromatin proteins include other DNA-binding proteins (e.g., Alba, HU) and non-nucleosomal histones (e.g., α3-truncated histone)^12,36^. In Archaea that are histone-poor or in which histones are absent, alternative chromatin proteins are abundant, whereas in the Loki and Hod Asgard archaeal groups, these alternative chromatin proteins are largely absent and histones are abundant, consistent with the ability of Hod histones to assemble onto DNA in vitro^32^ and suggesting a histone-based genome packaging as a feature of cultured Asgard archaea (Fig. 1a).

The histone genes in certain Asgard archaea are exceptionally diverse when compared with those of other Archaea, which typically encode only 1–3 histones, most belonging to a single canonical hypernucleosome-forming family ^21,37–40^. For example, cultured Loki species have 8–11 histone genes and Hod-HC1 has 6. Our phylogenetic, structural, and sequence analyses classified these histones into three families: untailed, canonical archaeal histones (HLoA in Loki and HMpA in Hod); tailed histones with lysine-rich N-terminal extensions (HLoB in Loki and HMpB in Hod), and family-specific, non-canonical variants (HLoC in Loki and HMpC in Hod; Fig. 1b; Supplementary Table 2). Each family contains histone isoforms reminiscent of the diversity of histone variants in eukaryotes.

To place these Asgard archaeal histones in the context of histone evolution, we mapped them onto a sequence similarity network of all known histone-fold proteins (Fig. 1c). Canonical archaeal and eukaryotic histones form distinct clusters, but the Asgard tailed histones, HLoB and HMpB, occupy an intermediate position within the archaeal cluster yet possess features of eukaryotic histones (disordered tails and increased lysine content) (Supplementary Fig. 3) similar to the highly conserved eukaryotic histone H4. This analysis also identified a previously unrecognized histone family, HLoC, which has the histone fold but occupies a peripheral position in the sequence similarity network, consistent with greater sequence divergence. HLoC histones are restricted to Loki and relative lineages (e.g., Helarchaeia, Sigynarchaeia) (Supplementary Table 3). They were excluded from further analyses owing to their highly divergent sequences and limited phylogenetic distribution.

Whereas most archaeal histones consist solely of a histone fold domain, the Asgard histones, HLoB and HMpB, also have disordered N-terminal tails. Proteomic and transcriptomic profiling of both Loki and Hod showed that these tailed histones are among the most abundant proteins in these cells (Fig. 1d). According to structural predictions, Asgard tailed histones should form nucleosome-like particles with their tails exposed on the surface, as in eukaryotic nucleosomes (Extended Data Fig. 4). Immunostaining of the enrichment culture of Loki-B35 with an antibody specific to tailed histones (Supplementary Fig. 4) showed that these tailed histones co-localize with DNA in the nucleoid of Loki cells (Fig. 1e; Supplementary Fig. 5).

We conclude that histone families diversified and became the dominant chromatin proteins in at least two major groups of Asgard archaea, present at levels compatible with genome-wide packaging and with tails prone to post-translational modification.

### Asgard archaeal histones have diverse post-translational modifications

Post-translational modifications of histones are a crucial feature of eukaryotic chromatin that enables genomic regulation. To examine whether similar histone modifications exist in Archaea, we used proteomics on acid-extracted histones from multiple lineages of archaea. To identify hPTMs systematically, we used a standard pipeline of histone enrichment, multi-protease digestion for maximal tail-peptide coverage, and PTM-aware database searching (Fig. 2a). Our taxon sampling included 15 species of Archaea, among which were three species of Asgard archaea (Loki-B35, Loki-B36, and Hod-HC1) and three archaeal reference species (*Methanospirillum stamsii*, *Methanobrevibacter cuticularis*, and *Thermococcus gammatolerans*)^31,41^, as well as two reference eukaryotes (*Saccharomyces cerevisiae* and *Arabidopsis thaliana*). Overall, among the archaeal histones abundant enough to yield sufficient peptide coverage for hPTMs, we detected lysine methylation (me) and acetylation (ac), which are common eukaryotic hPTMs (Fig. 2a). The histone-fold core of archaeal histones was also methylated on conserved acidic residues (glutamate (E) and aspartate (D); Extended Data Fig. 5). Acidic-residue methylation such as methylation of histone H4 on residue D24 (H4D24me1) and on residue E53 (H4E53me1) is also found in eukaryotic histones^42,43^, suggesting that these types of hPTMs may predate eukaryogenesis. We assessed whether these modifications contribute to nucleosome stacking through molecular dynamics (MD) simulations. However, MD simulations showed that hypernucleosome assemblies remained largely unchanged in the presence or absence of hPTMs, with only minimal effects under the tested conditions (Supplementary Fig. 6). We detected no lysine trimethylation, ubiquitination, or phosphorylation on any archaeal histones, suggesting that these hPTMs either occur below the detection limits of our approach or are absent in these Archaea.

**Figure 2.**
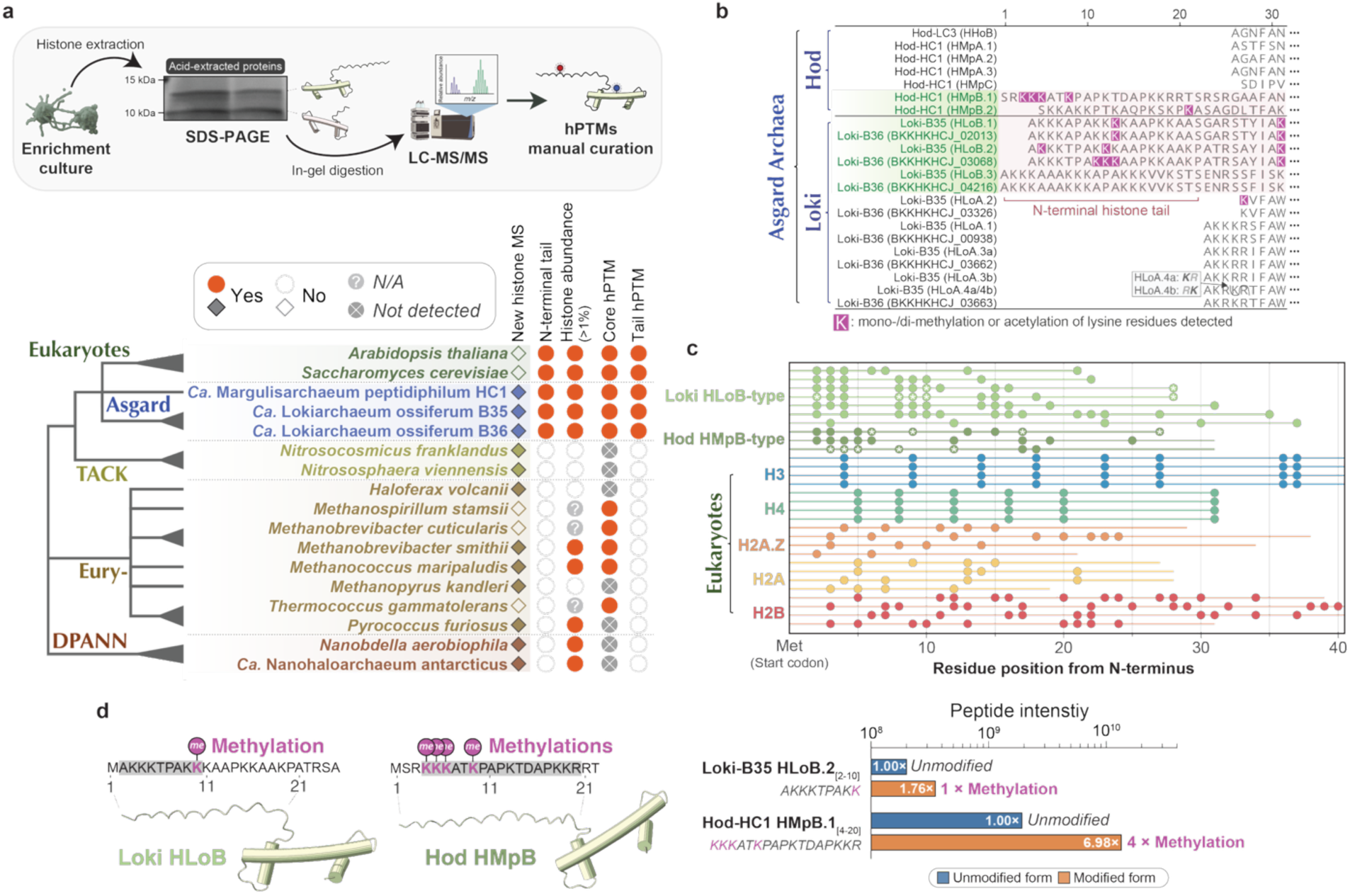
Asgard archaeal histones carry diverse post-translational modifications. **a**, hPTM landscape across the archaeal tree of life. Mass spectrometry workflow for cross-species hPTM detection: protein extraction, histone enrichment, multi-protease digestion, LC-MS/MS, and PTM-aware database search applied uniformly across multiple DPANN, Euryarchaeota, TACK, Asgard (Loki, Hod), and eukaryotic references. hPTMs were not reliably identified in some species (e.g., TACK, DPANN) due to insufficient histone enrichment, resulting in a peptide coverage too low for accurate PTM detection. **b**, Sequence alignment of histone tails showing site-specific modifications (acetylation and methylation) mapped to individual residues in Asgard archaeal histones. Modified lysine residues detected in Asgard archaea are colored purple. Comprehensive modification data (methylation and acetylation for each site) is provided in Extended Data Fig. 5. Boxed KR/RK motifs indicate the only sequence difference between HLoB.4a and HLoB.4b. **c**, Alignment of N-terminal tail residue positions from eukaryotic and Asgard tailed histones. Lysine positions across aligned histone tail sequences are shown as filled circles. Lysine positions detected as modified in Loki and Hod tailed histones are marked with white asterisks. **d,** Targeted peptide quantification of Asgard histone tail modifications. Left: structural representations of the Loki HLoB and Hod HMpB tailed histones with methylation sites indicated on the peptide sequences. Right: peptide intensity comparison showing enrichment of modified (methylated) tail peptides over their unmodified counterparts for Loki-B35 HLoB.2_[2–10]_ and Hod-HC1 HMpB.1_[4–20]_. Fold-change values are indicated on each bar. Assigned MS/MS spectra for detected modifications are shown in Extended Data Fig. 6. Residues are numbered from position 2, as the initiator methionine is absent in the mature polypeptide.

We discovered histone modifications of lysine residues in histone tails in the two independent Asgard archaeal groups (Loki and Hod) (Fig. 2b; Extended Data Fig. 6; Supplementary Fig. 7). In particular, we observed frequent methylation whereas tail acetylation was not detectable except in a single instance that may reflect a low abundance (Supplementary Fig. 7). Comparison of these histone tails with those of eukaryotes revealed lysine residues at comparable distances from the N-terminus with the potential to be modified as in eukaryotes (Fig. 2c). This positional correspondence across phylogenetically distinct lineages may suggest that they are subject to similar selective constraint, although the lack of sequence homology between tails precludes direct comparisons (Fig. 2c).

In eukaryotes, histone marks can be relatively abundant when compared to the unmodified residues^44–46^. Similarly, peptide quantification in Loki and Hod showed an enrichment of modified histone tail peptides relative to their unmodified counterparts. For example, modified forms of the Loki tailed histone peptide (HLoB.2_[2-10]_) and the Hod tailed histone peptide (HMpB.1_[4-20]_) both showed substantial enrichment over the corresponding unmodified peptides (Fig. 2d). These quantitative differences indicate that modified histone tail peptides are abundant in Asgard archaea and are likely not random noise. Collectively, the position-specific modification of tail lysines in two independent Asgard groups suggests these marks may be under selective constraint. We conclude that Asgard archaea evolved site-specific, enriched histone tail modifications.

### Asgard archaea have enzymes that write and erase histone tail acetylation

Regulation of gene expression by modifying chromatin requires enzyme writers and erasers that deposit hPTMs and remove hPTMs. To identify histone-modifying enzymes in the representative Asgard archaea Loki-B35, we combined multiple bioinformatic approaches comprising genomic survey, phylogenetics, and structure-based homology detection with AlphaFold-based predictions of protein–protein interactions to identify histone interactors (see Methods). These genomic surveys revealed that Asgard archaea with tailed histones (Loki and Hod) have an extensive repertoire of chromatin machinery, including writers, erasers, and histone proteins (Fig. 3a; Supplementary Table 4). Yet we identified no dedicated ‘reader’ domains, which bind modified histone tails in eukaryotes, nor a complete histone methylation regulatory pathway, consistent with a previous genomic study of chromatin machinery in Archaea^31^. By contrast, we found many acetylation writers (‘GCN5-related *N*-acetyltransferase’) and erasers (‘Histone deacetylase domain’ and ‘Sir2 family’). Therefore, we focused on characterizing these potential histone tail acetylation writers and erasers. In the Loki-B35 strain, we identified 63 candidate histone acetylation writers and 6 histone acetylation erasers (Supplementary Table 5).

**Figure 3.**
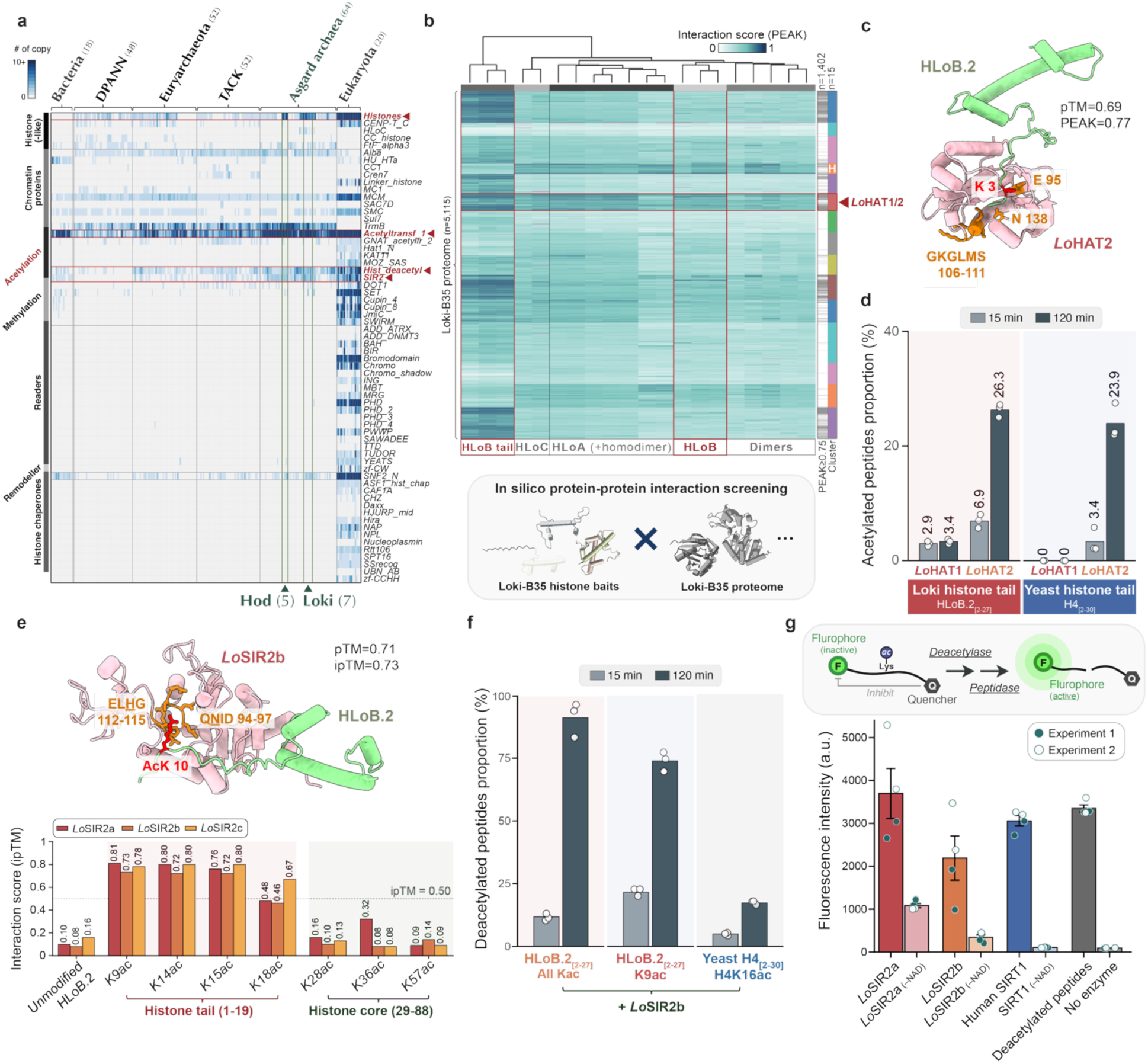
Identification and in vitro characterization of Asgard archaeal histone writers and erasers. **a**, Genomic distribution of chromatin-associated proteins across the tree of life (*n* = 255, see Supplementary Table 4). Key chromatin domain families were identified based on previous studies^12,20,31^. The heatmap shows the presence and copy number (0 to 10+, blue gradient) of chromatin-related protein families across representative genomes. Species are organized by major phylogenetic groups (left to right: Bacteria, DPANN, Euryarchaeota, TACK, Asgard archaea, Eukaryotes). Green arrows highlight Promethearchaeia/Lokiarchaeia (Loki) and Hodarchaeales (Hod), which show the most extensive chromatin protein repertoires in Archaea. Red arrows denote histone and acetylation-related chromatin modifiers characterized in this study. **b**, Systematic protein-protein interaction screening using AlphaFold2-multimer (>100,000 structural predictions). Loki histones in different forms (full-length, truncated tails, dimers; *n* = 21 baits) were screened against the Loki proteome (*n* = 5,115 prey). The heatmap shows interaction scores; magenta indicates failed predictions (*n* = 161), typically due to sequence length limitations or insufficient structural templates. Horizontal clusters (*n* = 5, grayscale) group baits by interaction patterns; vertical clusters (*n* = 15, colored) group prey proteins. The ‘H’ cluster contains 11 Loki histones (HLoA, HLoB, HLoC). Red arrowheads highlight the protein cluster (*n* = 250, Supplementary Table 5) enriched for high-confidence interactions with both tail-truncated and full-length HLoB histones, including 14 putative histone acetyltransferases (e.g., *Lo*HAT1/2) identified in the Loki-B35 proteome. **c**, AlphaFold2 structural prediction of the *Lo*HAT2–HLoB.2 complex. *Lo*HAT2 (pink) engages the N-terminal tail of HLoB.2 (green). Predicted catalytic interface residues on *Lo*HAT2 (orange: Gly106–Ser111 (GKGLMS), Glu95, Asn138) are related to acetyl-CoA binding sites and are positioned to contact the first accessible lysine on the histone tail (red: Lys3). Interacting residues were identified based on conserved active-site positions in a reference GNAT acetyltransferase family ^63^ (e.g., UniProt Q86UY6, P43577, P76112). A more detailed structural prediction is shown in Supplementary Fig. 8. **d**, In vitro HAT activity assays using synthetic Loki histone tail peptides (HLoB.2_[2-27]_) or yeast histone HHF1 tail (H4_[2-30]_) as substrates were incubated with Loki HAT enzymes (*Lo*HAT1, *Lo*HAT2). Acetylated peptide fractions (%) were quantified by mass spectrometry at 15 and 120 min after subtraction of no-enzyme controls. Mass spectrometry reveals that *Lo*HAT2 exhibits robust acetyltransferase activity, generating approximately a 10-fold higher acetylated peptide signal after 120 minutes compared to unincubated controls. **e**, AlphaFold3 predictions of *Lo*SIR2b–HLoB.2 complexes with an acetylated lysine residue. The structural model (top) shows *Lo*SIR2b (pink) engaging acetylated HLoB.2 (green, K9ac). Conserved sirtuin catalytic residues on LoSIR2b, including key NAD^+^-binding pocket (orange; Gln94–Asp97 (QNID) and proton acceptor (orange; Glu112–Gly115 (ELHG)), contact the acetylated lysine (red: AcK10). Bar graphs (bottom) quantify interaction confidence across specific acetylated lysine positions. Acetylation markedly enhances the predicted inter-protein interface confidence, suggesting that tail acetylation may promote *Lo*SIR2 substrate recognition. A more detailed structural prediction is shown in Extended Data Fig. 7. **f**, In vitro SIR2 deacetylation assays using synthetic Loki histone tail peptides. Fully acetylated HLoB.2_[2-27]_ (all Kac), mono-acetylated HLoB.2_[2-27]_K9ac, or yeast HHF1 histone tail (H4_[2-30]_K16ac) substrates were incubated with recombinant *Lo*SIR2b in the presence or absence of NAD^+^. Deacetylated peptide proportions (%) were quantified by mass spectrometry at 15 and 120 min after subtraction of no-enzyme controls. *Lo*SIR2b efficiently deacetylates Loki histone tail peptides and shows reduced but detectable activity on the yeast H4 substrate. Activity is abolished in the absence of NAD⁺, consistent with sirtuin-type catalysis (Supplementary Fig. 11). **g**, Fluorophore-based deacetylation assay measuring NAD^+^-dependent activity of recombinant Loki SIR2 proteins. The schematic (left) illustrates the assay principle: deacetylation of an acetylated lysine substrate by a sirtuin enables subsequent peptidase cleavage, releasing a fluorophore for quantification. The bar chart (right) shows endpoint fluorescence intensity (a.u.) after 60 min incubation at 20°C (excitation 355 nm, emission 460 nm). *Lo*SIR2a and *Lo*SIR2b (∼1.4 µM) display deacetylase activity comparable to human SIRT1 (∼1.0 µM; positive control). Activity is abolished in the absence of NAD^+^ (‘-NAD’), confirming NAD^+^-dependent catalysis. Additional controls include deacetylated peptides (pre-cleaved positive control) and no enzyme (substrate-only negative control). Individual data points from two independent experiments with duplicates are shown (*n* = 4). Error bars represent the standard error of the mean.

GCN5-related N-acetyltransferases (GNAT) are a large and functionally diverse enzyme family found across all domains of life, which can also acetylate non-histone substrates^47,48^. We therefore used AlphaFold-Multimer predictions to identify the subset with high-confidence interactions with histone tails. Genomic surveys and in silico screening identified candidate Asgard histone acetyltransferases (HATs) of the GNAT family as potential hPTM writers (Figs. 3a,b; Supplementary Table 6). Our predictions indicated that candidate HATs should interact with Asgard tailed histones, with high-confidence protein interactions localized to the histone tail region (Fig. 3c; Supplementary Fig. 8). To test the writer function, we expressed and purified recombinant *Lo*HAT1 and *Lo*HAT2 and incubated them with synthetic HLoB histone tail peptides in vitro. Mass spectrometry results showed that only *Lo*HAT2 had robust acetyltransferase activity, with an ∼10-fold higher signal for acetylated peptides compared to unmodified peptides after 120 minutes of incubation, identifying *Lo*HAT2 as a functional histone-tail acetyltransferase (Fig. 3d; Supplementary Fig. 9). *Lo*HAT2 also acetylated the yeast histone H4 N-terminal tail, consistent with a broad substrate specificity or a broadly conserved recognition of lysine-rich disordered tails^49^. We conclude that *Lo*HAT2 is a functional acetyltransferase able to acetylate lysine residues of Asgard histone tails.

Our genomic surveys also identified Asgard archaea that encode SIR2-family histone deacetylases, including three NAD-dependent histone deacetylase SIR2 proteins in Loki-B35 (e.g., *Lo*SIR2a–c). Asgard SIR2 proteins and other SIR2 proteins share the substrate recognition residues and the conserved catalytic core, including NAD^+^-binding residues (Supplementary Figure 10). Consistent with this, AlphaFold3 predicted that acetylated histone tails should bind *Lo*SIR2, whereas unmodified tails should not (Fig. 3e; Extended Data Fig. 7).

To test whether *Lo*SIR2 deacetylates histone tails directly, we performed in vitro deacetylation assays with recombinant *Lo*SIR2 with synthetic acetylated Loki histone tail peptides. Mass spectrometry showed that *Lo*SIR2b deacetylated both substrates in a NAD^+^-dependent manner, with activity abolished in the absence of NAD^+^, consistent with sirtuin-type catalysis (Fig. 3f; Supplementary Figure 11). We also performed in vitro deacetylation assays with recombinant *Lo*SIR2 and generic acetylated peptide substrates fused to a fluorophore (Fig. 3g; Supplementary Figure 12). This commercial substrate was chosen to enable direct quantitative comparison of *Lo*SIR2 with eukaryotic SIR2 under identical conditions. Fluorophore-based assays confirmed that *Lo*SIR2 and the human SIR2 protein (SIRT1) both deacetylated the same acetylated substrates in an NAD^+^-dependent manner, confirming the acetylation eraser properties of *Lo*SIR2.

Together, these data show that Asgard archaea, like eukaryotes, possess both writer and eraser activities able to acetylate and deacetylate lysine residues in histone tails.

### Asgard SIR2 erasers complement the loss of their eukaryotic homolog

The lysine residues of histones assembled in chromatin are constrained within a structural context that limits their accessibility to modification. The histones in our in vitro assays were not subject to this constraint. Therefore, to test whether Asgard SIR2 enzymes deacetylate histone tails in chromatin, we employed the *S. cerevisiae* Cre-Reported Altered States of Heterochromatin (CRASH) assay, used for studying histone acetylation-mediated silencing^50–52^. In yeast, Sir2 primarily removes acetyl groups from H4K16ac on nucleosomal substrates to establish transcriptional silencing at telomeres and silent mating-type loci (*HMLα*/*HMRa*). In the CRASH assay, deletion of *SIR2* gene causes complete derepression of Cre recombinase located within *HMLα*, triggering an irreversible conversion of a euchromatic reporter from RFP to GFP expression (Fig. 4a). At the population scale, the frequency of RFP-to-GFP conversion reflects the underlying de-repression frequency, thus quantifying both deacetylase catalytic activity and the ability to recognize acetylated lysine on histone tails within assembled nucleosomes^53,54^.

**Figure 4.**
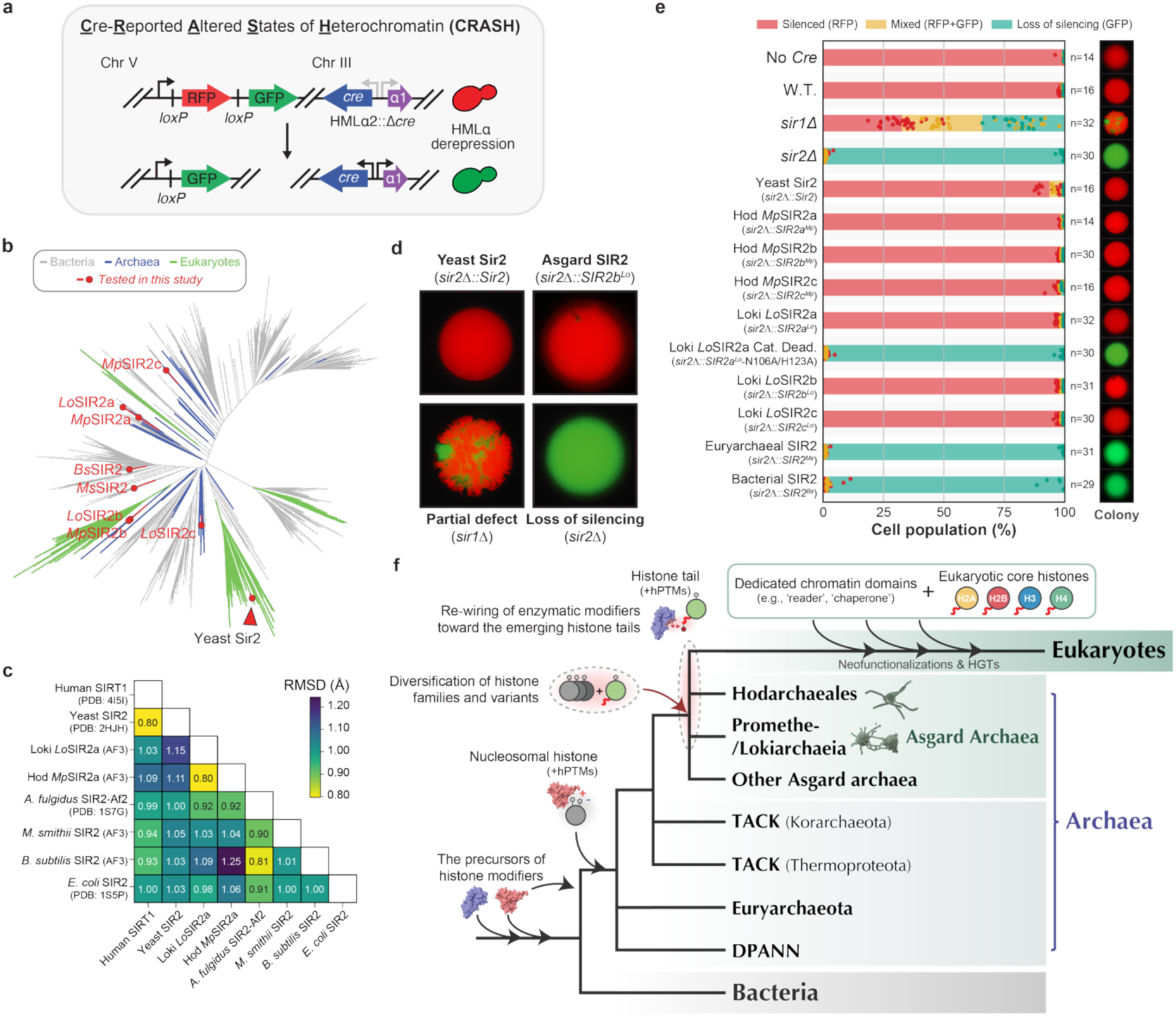
Asgard archaeal SIR2 enzymes functionally complement eukaryotic chromatin silencing. **a**, Yeast Cre-Reported Altered States of Heterochromatin (CRASH) assay design. Deletion of Sir2 abolishes H4K16 deacetylation and transcriptional silencing and expression of SIR2 homologs is scored for rescue of silencing. **b**, Phylogeny of the SIR2-family deacetylases. Asgard archaea encode at least three SIR2 paralogs: two clades (SIR2a, b) have shared ancestry between Loki and Hod, while SIR2c represents lineage-specific paralogs. Colored branches highlight eukaryotic (green), archaeal (blue), and bacterial (dark grey) clades. Red dots and nodes mark SIR2 proteins tested in CRASH complementation assays. Full phylogenies can be accessed at iTOL (https://itol.embl.de/tree/193171188528841782548812). **c**, Pairwise structural similarity of representative SIR2 proteins. Heatmap showing pruned Cα root-mean-square deviation (RMSD, Å) from pairwise structural alignments of the conserved sirtuin catalytic core (114 to 203 aligned Cα pairs after outlier pruning). Low RMSD values (typically <1.0 Å) across distant lineages highlight the strong conservation of the sirtuin catalytic fold. **d,** Representative colony fluorescence images. The control strain (functional Sir2) and *sir2Δ::SIR2cLo* (Asgard SIR2 rescue) show complete silencing (red), while *sir2Δ* shows a loss of silencing (green). *sir1Δ* shows intermediate silencing with sectored colonies. **e**, Quantification of CRASH assay results across all tested constructs (14-32 experimental replicates per strain). Stacked bars show cell population percentages: silenced (red, RFP), mixed (yellow, RFP+GFP), or loss of silencing (green, GFP). Constructs are listed along the x-axis. Loki and Hod SIR2 enzymes rescue silencing to near wild-type levels. Catalytically dead Loki SIR2 mutants, methanogen (Euryarchaeota) SIR2, and bacterial SIR2 do not complement silencing. **f**, Schematic model depicting the stepwise evolution of histone-based chromatin complexity from an ancestral archaeal state through Asgard archaea to eukaryotes. Histone-based packaging and sporadic hPTMs represent an ancestral archaeal baseline. Asgard lineages increased histone abundance, diversified tail-bearing variants, and rewired pre-existing modifying enzymes, creating a potential proto-epigenetic axis. Eukaryotic elaboration (including readers, complex targeting, four core eukaryotic histones) built upon this Asgard foundation.

Asgard archaea have distinctive SIR2 expansions, with Loki and Hod each encoding three paralogous clades (Fig. 4b). SIR2-family proteins have a complex evolutionary history^55^. Given this, to test Asgard SIR2s for their deacetylase activity in a living cell, we first selected representative homologs from both Asgard archaea and other clades without inferring ancestral relationships to eukaryotic enzymes (Fig. 4b). Whereas pairwise structural comparison indicates that the main catalytic domain is strongly conserved across lineages (Fig. 4c), the yeast Sir2 N-terminal domain (DUF592 domain) is a lineage-specific innovation that evolved within the Saccharomycotina to recruit Sir2 into a yeast-specific heterochromatin complex^56^. We took advantage of this yeast-specific feature to directly compare prokaryotic SIR2’s for their histone deacetylase activity. We fused the N-terminal DUF592 to each prokaryotic SIR2 homolog, completely replacing the endogenous *SIR2* gene with these chimeras to enable their recruitment into the SIR complex. We then tested whether Asgard SIR2 enzymes would complement endogenous Sir2 and silence Cre (Fig. 4a). Expression of Loki and Hod SIR2 enzymes in the absence of endogenous yeast Sir2 silenced Cre at levels similar to WT cells (Figs. 4d, e). Importantly, the catalytically dead Loki SIR2 mutant (*Lo*SIR2a-N106A/H123A) failed to complement, confirming that rescue depends on the deacetylase activity of Asgard SIR2s.

Sir2 homologs from methanogenic Euryarchaeota (*Methanobrevibacter smithii*) and Bacteria (*Bacillus subtilis*) failed to rescue yeast *sir2Δ* mutants (Fig. 4e), suggesting that these non-Asgard prokaryotic enzymes are incompatible with the assembly or function of the yeast SIR (Silent Information Regulator) complex. Altogether, these data show that, unlike the tested bacterial and euryarchaeal homologs, Asgard SIR2 enzymes are functionally compatible with eukaryotic chromatin and can deacetylate histone tails in the context of the nucleosome, pointing to a pre-eukaryotic molecular foundation for chromatin regulation.

## Discussion

In this study, we showed that Asgard archaea—the closest prokaryotic relatives of eukaryotes—have complex histone-based chromatin and the regulatory machinery required to write and erase hPTMs. We report that several key molecular components of eukaryotic chromatin regulation were already present in two phylogenetically independent Asgard lineages, Loki and Hod, before the emergence of eukaryotes. Like eukaryotes and unlike other Archaea, these Asgard archaea possess multiple families of histone variants and tailed histones expressed at high levels, which is compatible with their role as major components of chromatin.

Lysine residues on histone tails can be acetylated by HAT writers and removed by SIR2 erasers like their eukaryotic counterparts. Asgard SIR2 erasers act on their substrates in eukaryotic chromatin in vivo, whereas some tested bacterial and euryarchaeal SIR2 homologs do not, indicating that compatibility with eukaryotic chromatin is not a ubiquitous property of SIR2-family deacetylases. This result demonstrates that Asgard SIR2 enzymes can recognize and deacetylate histone tails, establishing a continuity of molecular mechanisms between Asgard and eukaryotic chromatin. Consistent with active deacetylation, our mass spectrometry data show that histone tails are predominantly methylated. The hardly detectable level of histone tail acetylation suggests that SIR2 erasers maintain low steady-state levels of tail acetylation while core-residue acetylation, which is not a SIR2 substrate, remains readily detectable. Although we identified histone tail methylation, we found no orthologs of SET domain eukaryotic histone methyltransferases or Jumonji domain demethylases in our representative Asgard archaea. Further, homologs of eukaryotic reader domains with the capacity to bind acetylated or methylated histone tails were not identified, suggesting that the histone methylation system evolved independently in the Asgard lineage.

The absence of identifiable reader domains for both acetylation and methylation raises the question of how these modifications exert their effects. Asgard hPTMs could regulate chromatin through intrinsic biophysical effects. Charge-based alterations and conformational changes at modified residues could directly modulate interactions without requiring dedicated recognition proteins^57^, as demonstrated for tail hPTM effects on nucleosome electrostatic potentials^58^. Moreover, because histone acetylation depends on acetyl-CoA availability, such a system could couple chromatin state to cellular metabolism, potentially through promiscuous recognition by proteins capable of acetyl-lysine binding, as observed in bacterial transcription factors^59–61^. Determining the function of histone modifications will require the development of genetic tools for Asgard archaea, and both regulatory and non-regulatory interpretations of these hPTMs remain open.

Based on our findings, we hypothesize a stepwise model for the emergence of histone-based chromatin regulation (Fig. 4f). In this model, genome packaging mediated by untailed histones was broadly distributed in ancestral Archaea. In Asgard archaea, histones became more abundant and diversified to include a family of tail-bearing histone variants, establishing a dominant, modification-competent substrate. With the addition and rewiring of writer and eraser enzymes, the capacity for histone-based genome regulation was thus accessible to Archaea before eukaryogenesis. The subsequent integration of reader domains, complex targeting mechanisms, and the stabilization of the four core histones by chaperones consolidated this pre-existing capacity into the genome regulation by chromatin that underlies the complexity of eukaryotes.

## Figures

**Extended Data Figure 1.**
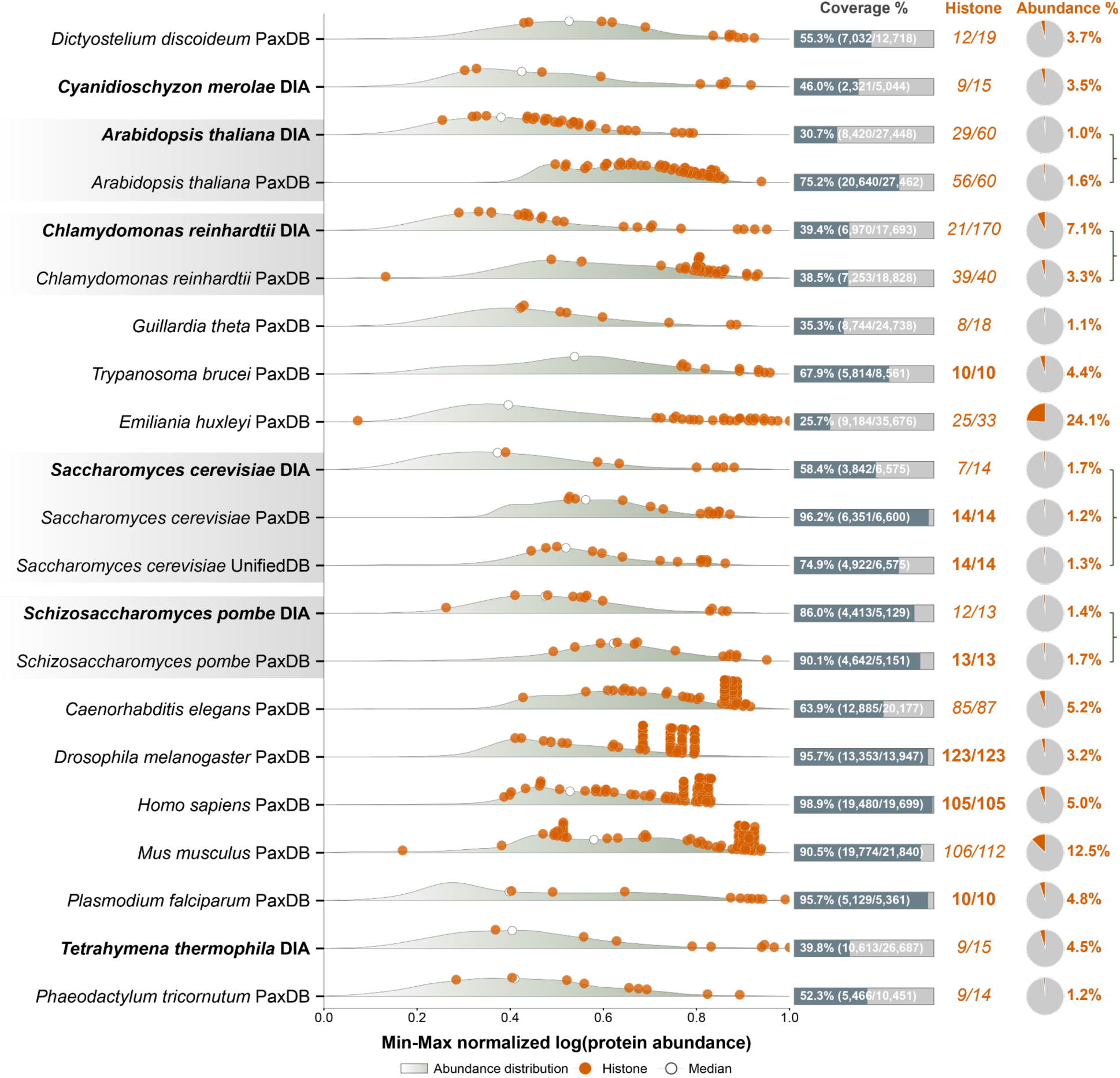
Eukaryotic histone abundance establishes a lower bound for genome-packaging levels of histones. Ridge plots show how protein abundances are distributed across eukaryotic proteomes. These abundances are measured using intensity-based absolute quantification (iBAQ) or parts per million (ppm). The protein abundances are displayed as min-max normalized values on a log scale, scaled from 0 to 1 within each proteome. Each ridge represents the complete abundance profile of a single organism. Histones are highlighted in red, with the median normalized histone abundance indicated by an open circle. Right-hand columns provide: (i) ‘Coverage %’, percentage of the predicted proteome detected by mass spectrometry; (ii) ‘Histone’, number of histone paralogs detected out of the total predicted histone genes (detected/predicted); (iii) ‘Abundance %’, percentage of total cellular protein represented by histones, calculated as the sum of histone protein abundances divided by the sum of all protein abundances. Histone abundance ranges from 1.0% (*Arabidopsis thaliana*) to 24.1% (*Emiliania huxleyi*) across diverse eukaryotic lineages. All eukaryotic species exceed the ∼1% threshold established for genome-packaging levels of histones (see Fig. 1a), validating this benchmark for genome-packaging function. Data sources: whole-cell proteomics generated in this study (*n* = 6; indicated as ‘DIA’, bold text) and reference datasets from PaxDb (‘PaxDB’)^64^ and the unified protein abundance dataset (‘UnifiedDB’)^34^. Each database may use different reference proteomes; the number of detected histone paralogs for the same species may vary between sources.

**Extended Data Figure 2.**
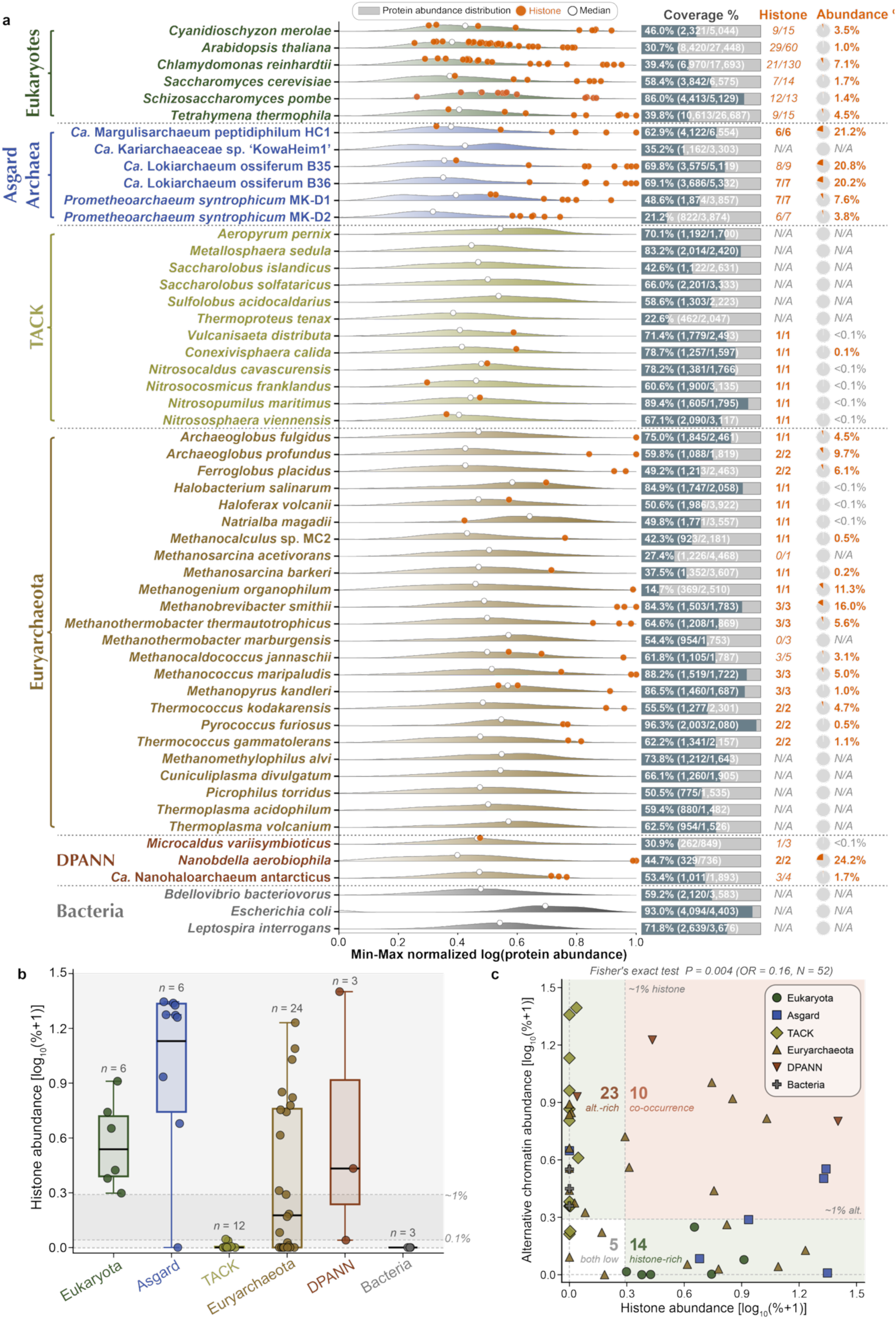
A tree-of-life proteome analysis reveals the evolution of histone abundance in Archaea. **a**, Ridge plots showing the distribution of normalized protein abundances across the proteomes of representative organisms spanning major domains of life, including eukaryotes, Asgard archaea, TACK, Euryarchaeota, DPANN, and Bacteria. Min-max normalization scales each proteome independently: the lowest-abundance protein is set to 0 and the highest-abundance protein to 1, enabling direct comparison of relative abundance distributions across species with different dynamic ranges. Each distribution reflects the proteome-wide normalized protein abundance profile of a single organism, with proteome coverage (fraction of detected proteins) indicated as ‘Coverage %’. Nucleosomal histones detected in each organism are labeled and highlighted (orange circles), and the median normalized protein abundance value is shown (white circle). The percentage of total protein abundance (rounded to one decimal place) attributable to histones is indicated for each organism. **b**, Distribution of histone protein abundance across phylogenetic groups. Boxplots with overlaid points showing histone protein abundance for each taxonomic group in log_10_(%+1)-transformed representation. Each point represents a single species, colored by taxonomic affiliation. Boxes indicate the interquartile range (IQR) with the median marked as a horizontal line, and whiskers extend to 1.5× IQR. Sample sizes (*n*) are indicated above each group. **c**, Mutual exclusivity between histone and alternative chromatin protein abundance. Scatterplot showing the relationship between histone protein abundance (x-axis) and the summed abundance of alternative chromatin-associated proteins (y-axis) across 54 species, with values log_10_(%+1)-transformed. Two species (*M. marburgensis*, *M. alvi*) with no detected chromatin proteins were excluded. Each point represents a single species, colored by taxonomic affiliation. Species are classified into four quadrants based on a 1% abundance threshold (dashed lines): alt.-rich (low histone, high alternative), co-occurrence (both high), both low, and histone-rich (high histone, low alternative). Quadrant counts are shown in bold. Fisher’s exact test on the resulting 2×2 contingency table confirms a statistically significant pattern of mutual exclusivity (odds ratio = 0.16, *P* = 0.004, *n* = 52). An odds ratio below 1 indicates that species with a high abundance of one protein class are less likely to also have a high abundance of the other.

**Extended Data Figure 3.**
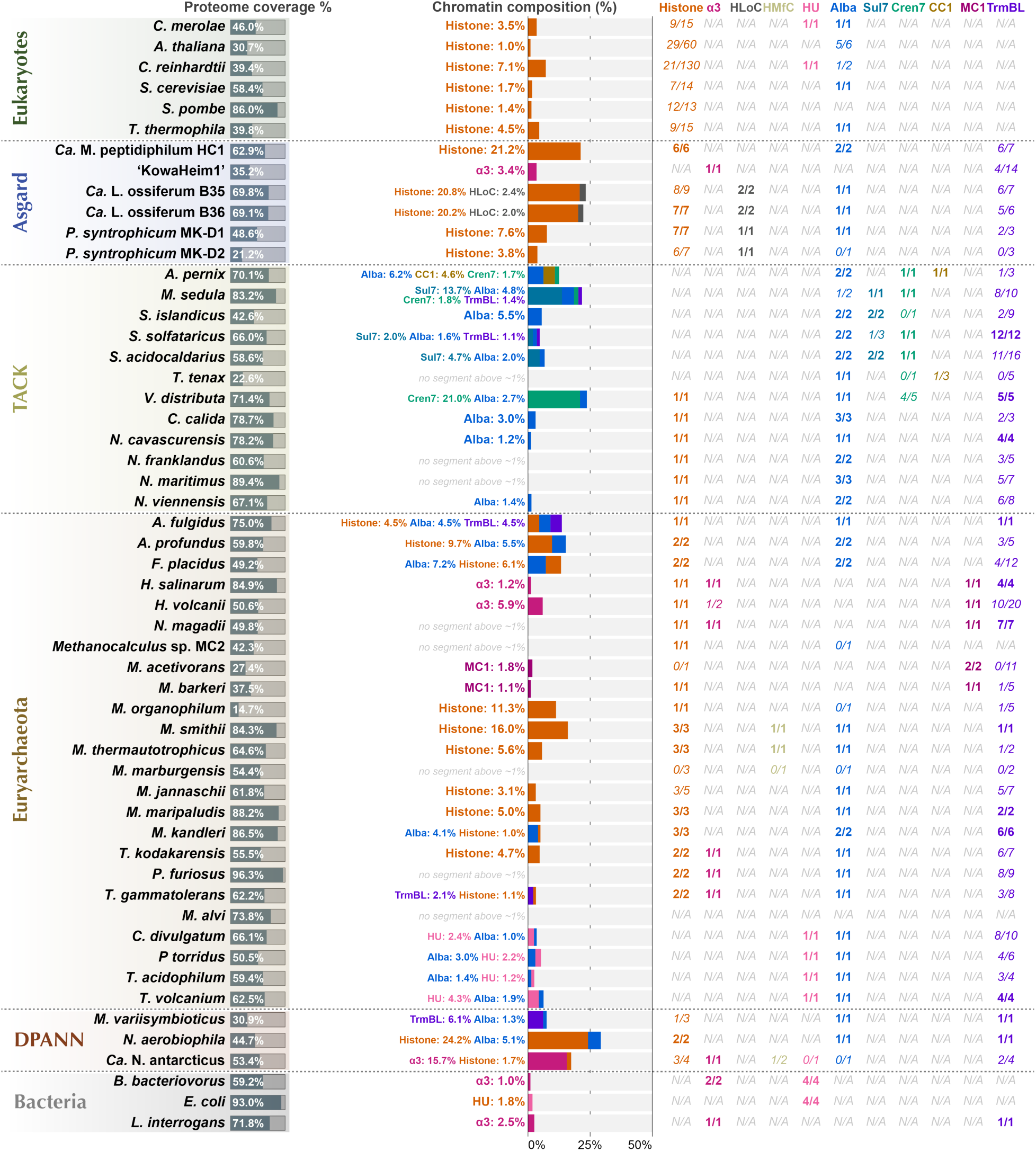
Chromatin protein composition across the tree of life. Stacked bar charts display the percentage composition of chromatin-associated protein families within each organism’s proteome, calculated as the sum of family-specific protein abundances divided by total proteome abundance. Protein families are color-coded: Histone (orange), α3-truncated histone (red), HLoC histone (face-to-face histone; light blue), HMfC histone (coiled-coil histone; dark blue), HU (pink), Alba (green), Sul7 (dark green), Cren7 (olive), CC1 (brown), MC1 (purple), and TrmBL (magenta). Individual family contributions of above ∼1% are labeled on each bar. Organisms are grouped by taxonomic domain (Eukaryota, Asgard archaea, TACK archaea, Euryarchaeota, DPANN archaea, Bacteria) and are separated by horizontal lines. The ‘Coverage %’ column (left) indicates the percentage of the predicted proteome that was detected by mass spectrometry. The right columns report the number of detected paralogs out of total genes (detected/total) for each chromatin protein family; ‘*N/A*’ indicates that the protein family is not encoded in the genome.

**Extended Data Figure 4.**
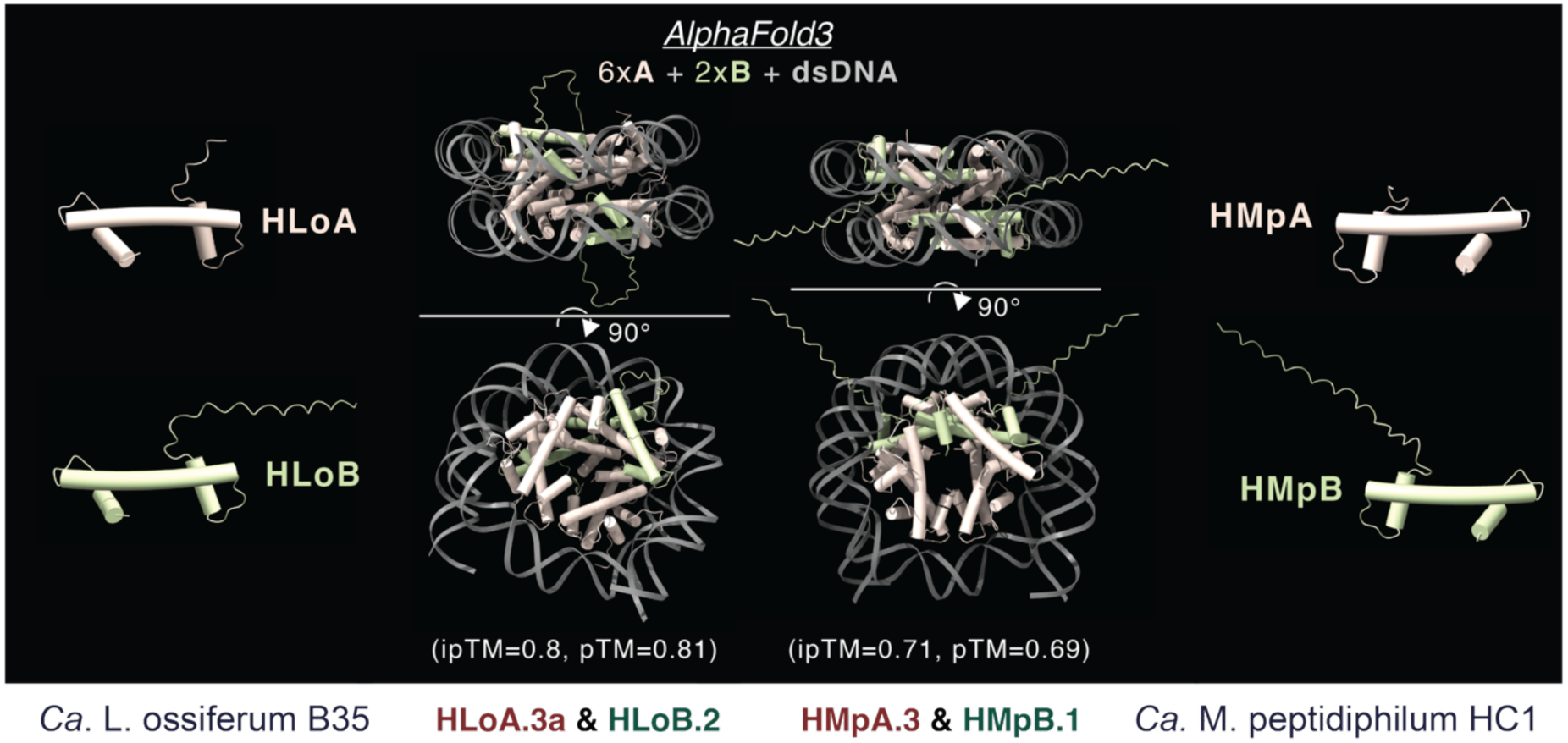
AlphaFold3 structural predictions of Asgard archaeal nucleosomes using tailed histones. AlphaFold3 predictions of nucleosome-like structures formed by Asgard archaeal histones with double-stranded DNA. Left: *Ca.* Lokiarchaeum ossiferum B35 nucleosome model (6×HLoA.3a + 2×HLoB.2 + dsDNA; ipTM=0.8, pTM=0.81). Right: *Ca.* Margulisarchaeum peptidiphilum HC1 nucleosome model (6×HMpA.3 + 2×HMpB.1 + dsDNA; ipTM=0.71, pTM=0.69). We show the highest-confidence assemblies after testing multiple stoichiometric combinations. The top panels show the individual histone subunit structures (HLoA, HLoB, HLoC for Loki; HMpA, HMpB, HMpC for Hod). Bottom panels display two views of the assembled nucleosome-like complexes, with DNA (gray) wrapped around the histone core.

**Extended Data Figure 5.**
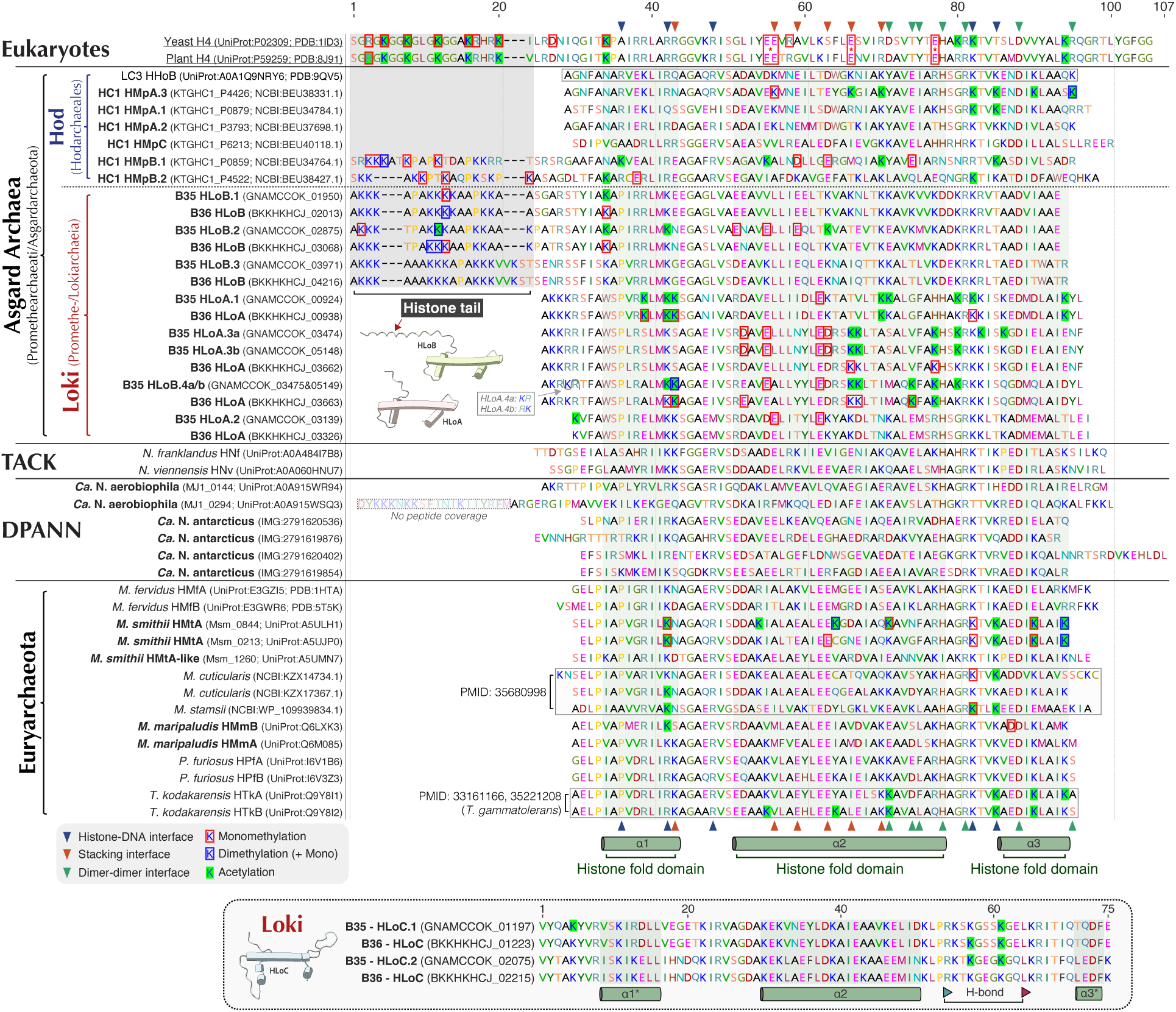
Histone post-translational modifications and sequence conservation across the tree of life. The figure shows multiple sequence alignments of histone and histone-like proteins from eukaryotes, Asgard archaea, TACK, DPANN, and Euryarchaeota, and it also includes annotated structural and functional features of the histone fold domain. Residues are shown starting from the position +1 after the initial M residue. Sequences are numbered according to the reference histone fold structure, with α-helices indicated. Conserved residues mediating histone-DNA contacts, stacking interactions, and dimer-dimer interfaces are annotated. The alignment shows known and newly identified histone post-translational modifications (hPTMs), including monomethylation, dimethylation, and acetylation at specific lysine residues. Structural schematics illustrate the distinct domain architectures of Asgard archaeal histone variants (HLoA, HLoB, HLoC). Annotated eukaryotic histone modifications are based on previously published data^42,65–67^.

**Extended Data Figure 6.**
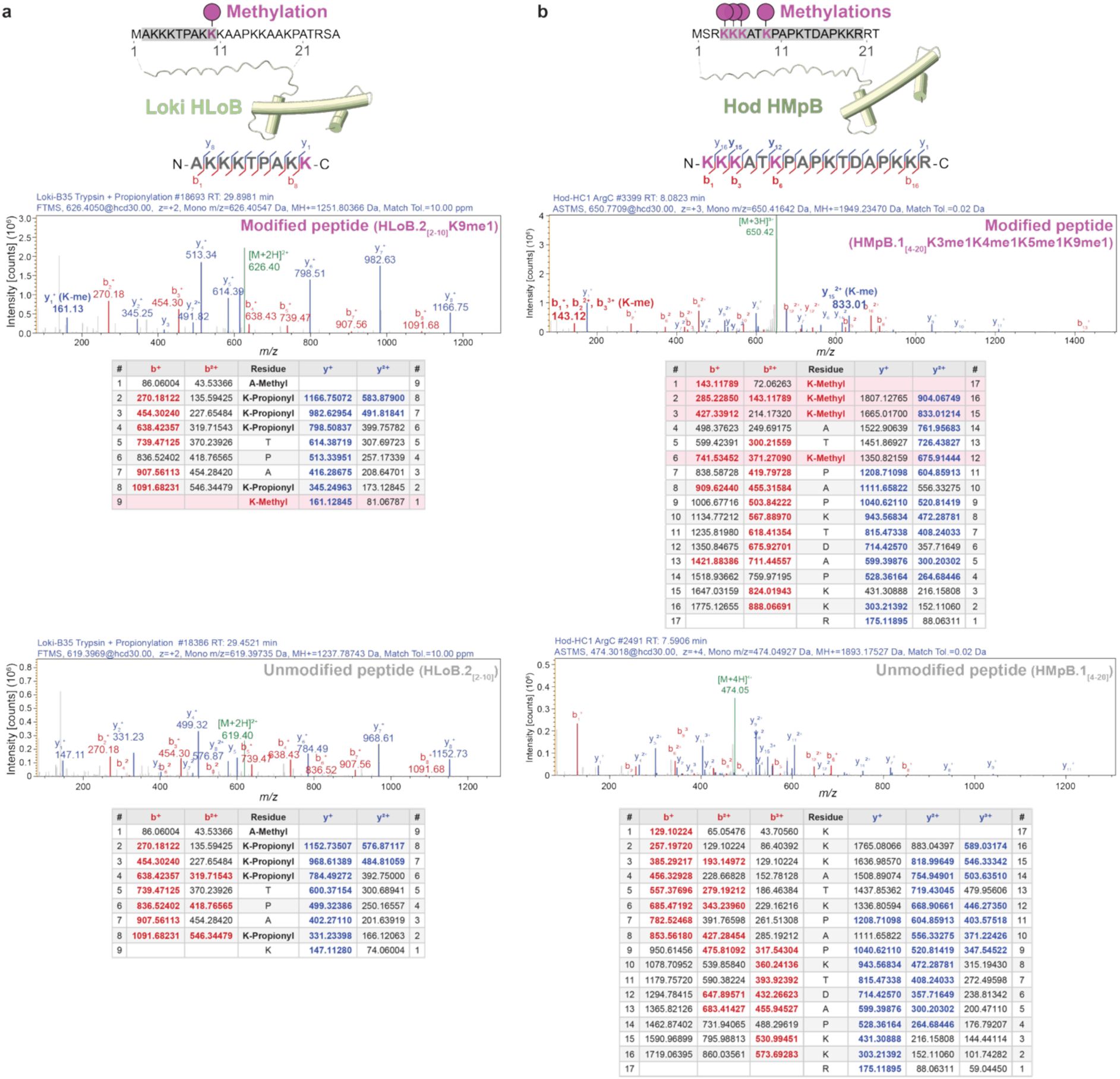
MS/MS validation of histone tail post-translational modifications in Asgard archaea. **a**, MS/MS spectrum and fragment ion table for the Loki-B35 HLoB tailed histone peptide AKKKTPAKK showing monomethylation at K9 (HLoB.2_[2–10]_K9me1). Histone residues are numbered canonically from the second residue, as the initiator methionine is absent in the mature polypeptide. Propionylation (pr) of unmodified lysine residues (K2, K3, K4, K8) results from chemical derivatization during sample preparation. The diagnostic y_1_^+^ ion localizes the modification to K9. **b,** MS/MS spectrum and fragment ion table for the Hod-HC1 HMpB tailed histone peptide KKKATKPAPKTDAPKKR showing methylation at K1, K2, K3, K6 residues of peptide (HMpB.1_[4–20]_K3me1K4me1K5me1K9me1). In both panels, spectra and fragment ion tables for the modified form are shown on top and those for the unmodified form on the bottom. Fragment ion abundance is plotted against mass-to-charge ratio (*m*/*z*). The *b* and *y* ions and their neutral losses of H_2_O are marked in red and blue, respectively, precursor ions in green, and unassigned peaks in light grey. Non-essential ion labels have been omitted for readability. Fragment ion tables list theoretical *m*/*z* values for detected ion series. pr, propionylation; me, methylation.

**Extended Data Figure 7.**
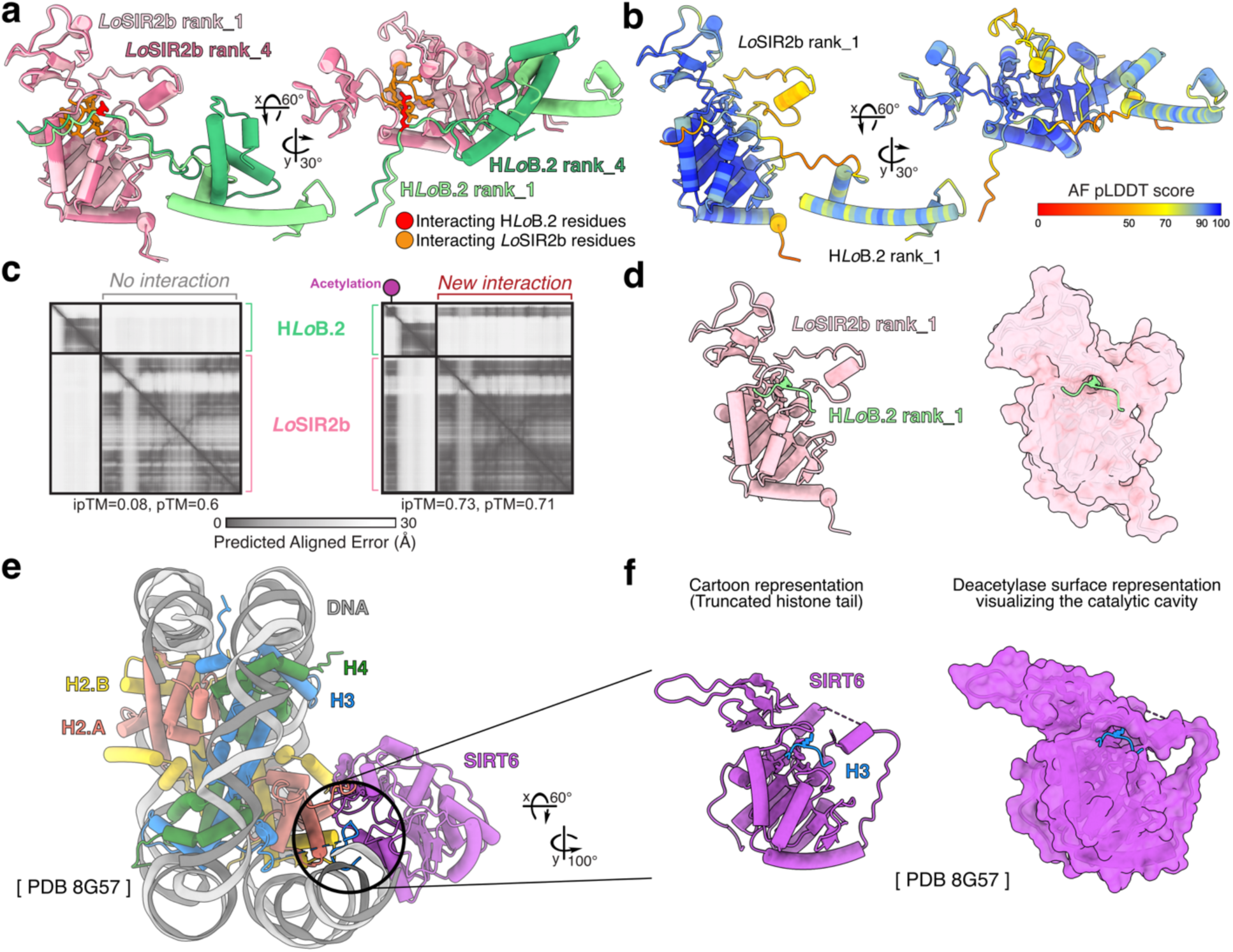
AlphaFold structural prediction of Asgard histone eraser. **a-d**, AlphaFold3 (AF3) predictions of *Lo*SIR2b–HLoB.2 complexes. **a**, Superimposed AF3 structural models of two most distinct prediction models of *Lo*SIR2b (two shades of pink) engaging acetylated HLoB.2 (two shades of green, K9ac) (rank 1 and rank 4). This panel illustrates the high prediction confidence in the protein core fold of both *Lo*SIR2b and HLoB.2 together with their interaction interface. The relative position of the histone is variable, due to the highly flexible histone tail. As displayed also in Figure 3e, conserved sirtuin catalytic residues on *Lo*SIR2b, including key NAD^+^-binding pocket (orange; Gln94–Asp97 (QNID)) and proton acceptor (orange; Glu112–Gly115 (ELHG)), contact the acetylated lysine (red: AcK10). Models shown in two orientations. **b**, Highest-ranked models of *Lo*SIR2b and HLoB.2 (rank_1), colored by the Predicted Local Distance Difference Test (pLDDT) score from AF3 and shown in two orientations. **c**, Predicted Aligned Error (PAE) plots from AF3 showing the *Lo*SIR2b–HLoB.2 interaction prediction difference with (left) and without (right) acetylation of K9ac. **d**, Highest rank models of *Lo*Sir2b and a truncated HloB.2 (rank_1), showing the histone tail interacting residue (AcK10), together with two flanking residues. Cartoon representation (left) and surface representation of *Lo*SIR2b (right) visualize the predicted catalytic cavity, where the histone tail is binding. **e,f**, Structure of the experimentally determined human Sirtuin 6 (SIRT6 in purple)—nucleosome complex (PDB: 8G57) displaying an interaction between SIRT6 and one of the histone H3 tails (blue). **f**, Magnified view of the model of SIRT6 and truncated H3 showing the histone tail interacting residue (Lys9), together with two flanking residues. The cartoon representation (left) and surface representation of SIRT6 (right) visualize the catalytic cavity, where the histone tail is binding, similarly as observed on the predicted *Lo*SIR2b model.

## Methods

### Whole-cell lysate proteomics sample preparation

Archaeal and eukaryote cultures were cultured under standard conditions appropriate to each lineage (Supplementary Table 7). Most of the samples were harvested during the mid-exponential phase by centrifugation, washed with anaerobic PBS, and flash-frozen in liquid nitrogen. Reference datasets were obtained from public repositories (e.g., PaxDB)^64^ or generated as indicated in Supplementary Table 8.

Cell pellets from archaeal cultures (typically 100 mg wet mass) were resuspended in 500 μl of Archaea Lysis (AL) buffer, which contained 50 mM Tris-HCl pH 7.4, 150 mM NaCl, 10 mM MgCl_2_, 1 mM EDTA, 0.1% IGEPAL CA-630, and an optional 0.1% Triton X-100, and was freshly supplemented with 1 mM DTT and 1× protease inhibitor cocktail (cOmplete, Roche). Cells were lysed by sonication (Branson Sonifier, microtip 220-B, 20% amplitude, 1 s on / 3 s off, 3 min total pulse time) on ice, with the temperature monitored to prevent overheating. Lysates were clarified by centrifugation at maximum speed (>16,000 g) for 20 min at 4°C. Supernatants were collected, and protein concentrations were estimated by absorbance at 280 nm (NanoDrop) using lysis buffer as a blank. Aliquots of 20–50 μl (depending on concentration) were submitted to the Vienna BioCenter Proteomics Facility for LC-MS/MS analysis. Remaining lysate was snap-frozen in liquid nitrogen and stored at -70°C. Alternative lysis protocols (e.g., phenol-chloroform, SDS-based) were used for specific lineages as detailed in Supplementary Table 7. Cell lysates were mixed with 2x S-Trap lysis buffer and prepared and tryptically digested for MS analysis using the S-Trap protocol (Universal MS sample Prep Kit with micro columns, ProtiFi) according to the manufacturer’s protocol.

### Acidic extraction of histone proteins from archaeal species

Histone-enriched fractions were prepared by acid extraction adapted for archaeal cells (Supplementary Tables 7, 9). Cell pellets (typically 100 mg wet mass) were centrifuged at 2,500 rcf for 10 min at 4°C and washed three times with 2 ml ice-cold Nuclei Isolation Buffer (NIB: 0.25 M sucrose, 60 mM KCl, 15 mM NaCl, 5 mM MgCl_2_, 1 mM CaCl_2_, 15 mM PIPES pH 6.8, 0.8% Triton X-100) without protease inhibitors, ZnSO_4_, or trichostatin A (TSA). Cells were then resuspended in 500 μl NIB supplemented with 2 ng/μl TSA, 2 mM ZnSO_4_, and 1× protease inhibitor cocktail. The entire suspension was transferred to 2 ml tubes containing 0.5 mm Disruptor Beads (cat. SI-BG05, Scientific Industries) filled until beads reached 1 mm below the liquid surface. Cells were lysed in a Precellys Evolution homogenizer (Bertin Technologies) at 8,000 rpm for 1 min, followed by a 30 s pause on ice, repeated three times. Lysates were collected into a fresh tube and centrifuged at maximum speed for 10 min at 4°C. The pellet was resuspended in 800 μl of 0.44 N H_2_SO_4_ and incubated for at least 1 h on ice. After centrifugation at 21,000 *g* for 10 min at 4°C, the acid-soluble supernatant was collected. The H_2_SO_4_ extraction was repeated on the pellet. Proteins were precipitated by dropwise addition of 270 μl of 6.1 N trichloroacetic acid (TCA) to each tube, vortexed, and incubated on ice for at least 30 min. Precipitated proteins were collected by centrifugation at maximum speed for 30 min at 4°C, washed twice with 500 μl cold acetone (centrifugation at maximum speed for 15 min between washes), and air-dried briefly. The final pellet was resuspended in 40–100 μl of 1× Laemmli buffer in 0.3× PBS (with 1 μl of 1 M Tris pH 8.0 per 20 μl if needed to adjust pH) and boiled at 95°C for 5 min. The extracted histone proteins were loaded on 15% precast SDS-gels (Cat. #456-1094, Mini-PROTEAN® TGX™ Precast Gels, BIO-RAD, USA) and were then stained using silver staining^68^ or Coomassie staining with MBS Coomassie Blue from the VBCF facility.

### In gel digest

Coomassie-stained gel bands were cut to 2–3 mm pieces, transferred to 0.6 ml tubes and incubated with different solutions by shaking for 10 min at room temperature followed by removal of the supernatant as follows: Gel pieces were washed with 200 µl 100 mM ammonium bicarbonate (ABC), destained by 2 repeated rounds of shrinking in 200 µl 50% acetonitrile (ACN) in 50 mM ABC and reswelling in 200 µl 100 mM ABC. Gel pieces were shrunk with 100 µl ACN before being reduced with 100 µl 10 mM dithiothreitol (DTT) in 100 mM ABC by incubation at 56°C for 30 min. After another shrinking step in 100% ACN they were alkylated with 100 µl of 25 mM iodoacetamide (IAA) in 100 mM ABC by incubation at RT for 30 min in the dark.

Wash steps were repeated as described for destaining and gel pieces were briefly dried by vacuum centrifugation after the final shrinking step. Gel pieces were soaked in 12.5 ng/µl trypsin (Trypsin Gold, Promega), Glu-C or chymotrypsin (both sequencing grade, Roche) in 100 mM ABC for 5 min at 4°C. Excess solution was removed, ABC was added to cover the pieces, and tryptic and Glu-C digest samples were kept overnight at 37°C. Chymotryptic digests were incubated at 25°C for 5 hr. In some cases, proteins were digested with Arg-C Ultra (Promega) using buffer and digest conditions recommended by the manufacturer. The supernatant containing generated peptides was transferred to a fresh tube and gel pieces containing non-propionylated peptides were extracted by addition of 20 µl 5% formic acid and sonication for 10 min in a cooled ultrasonic bath. This step was performed twice. All supernatants were unified.

If propionylation was applied, proteins were not reduced and alkylated but propionylated on the lysine side chains according to a protocol from the Bonaldi lab^69^ by swelling the gel pieces in a solution consisting of 15 µl propionic anhydride (#Nr), 26 µl 1 M ABC and 6 µl of 1 M sodium-propionate (Sigma). After shaking for 10 min at 37°C at 350 rpm, 80 µl 1 M ABC were added and the bands were incubated for 4 hr at 37°C at 1,400 rpm. Gel pieces were washed once with water and 2 times with 500mM ABC before being shrunk in 50% ACN, followed by another step in 100% ACN and a short drying step by vacuum centrifugation.

Gel pieces were washed once with water and 2 times with 500 mM ABC before being shrunk in 50% ACN, followed by another step in 100% ACN and a brief drying step by vacuum centrifugation. Proteins were digested with trypsin as described above. In some cases, propionylation was only performed on protein level and peptide extraction was done as for unmodified peptides described above.

In other cases, a second propionylation step was performed on peptide level: Propionylated peptides were extracted from the gel by addition of 100% ACN, extracts were dried down to below 5 µl by vacuum centrifugation and filled up to 15 µl with H₂O, supplemented with 2 µl of 1 M triethylammonium bicarbonate pH 8.5 (TEAB, Sigma) and 3 µl of a freshly mixed propionic anhydride (ratio 1:100 of propionic anhydride in 100 mM TEAB). After incubation for 2 min the sample was again vacuum-centrifuged to reach 1–5 µl and the propionylation was repeated a second time, before being quenched with 2 µl of 80 mM hydroxylamine. Samples were incubated for 20 min at 37°C before being acidified with formic acid. Gel digests were desalted using an Oasis HLB 96-well µElution Plate (with 2 mg Sorbent per Well, Waters) according to the manufacturer’s description. A similar aliquot of each digest was analyzed by LC-MS/MS.

### nanoLC-MS/MS Analysis

The nano HPLC system (Vanquish Neo UHPLC-System, Thermo Fisher Scientific) was coupled to an Orbitrap Exploris 480 or an Astral mass spectrometer equipped with a Nanospray Flex ion source (all parts Thermo Fisher Scientific) or a TimsTOF HT mass spectrometer (Bruker Daltonics). Peptides were loaded onto a trap column (PepMap Acclaim C18, 5 mm × 300 μm ID, 5 μm particle size, 100 Å pore size, Thermo Scientific) at a flow rate of 25 μl/min using 0.1% TFA as mobile phase.

After 5 min, the trap column was switched in line with the analytical column (PepMap Acclaim C18, 500 mm × 75 μm ID, 2 μm particles, 100 Å, Thermo Scientific operated at 30°C or Aurora Ultimate C18 25 cm × 75 μm ID, 1.7 μm particles, 120 Å, Ionopticks operated at 50°C). Peptides were eluted using a flow rate of 230 nl/min (PepMap) or 300 nl/min (Aurora), starting with the mobile phases 98% A (0.1% formic acid in water) and 2% B (80% acetonitrile, 0.1% formic acid) and linearly increasing to 35% B (or 45% B for propionylated peptides) over the next 60 min (or 35 min for in vitro acetylation assays), followed by a steep increase to 95% B and a 4-min hold at 95% B, and re-equilibration with 2% B for three column volumes (equilibration factor of 3.0).

For the analysis of whole cell digests, the Orbitrap Astral was operated in data-independent mode, performing a full scan in the Orbitrap every 0.6 s (m/z range 380–980 m/z, resolution 240,000, AGC target 1,000,000, maximum injection time 5 ms). MS/MS spectra were acquired in the Astral analyzer by isolating 5 Da windows across 380–980 m/z (resulting in 119 scan events per cycle) and fragmenting precursor ions with HCD collision energy of 25% with a maximum injection time of 3 ms or until an AGC target of 30,000 was reached. Fragment ions ranging from 150–2,000 m/z were acquired.

Several whole cell digests were analyzed on a TIMS quadrupole time-of-flight mass spectrometer (timsTOF HT, Bruker Daltonics) in DIA-PASEF mode, where 1 MS1 scan was followed by 8 DIA-PASEF frames. The CaptiveSpray source parameters were: 1,600 V capillary voltage, 3.0 L/min dry gas, and 180°C dry temperature. MS data was acquired in the MS scan mode using positive polarity, 100–1,700 m/z range. Mobility range was set to 0.64–1.42 V·s/cm², ramp time was set to 166 ms, and estimated cycle time was 1.52 s. Collision energy was 20 eV at 1/k₀ 0.6 V·s/cm² and 80 eV at 1/k₀ 1.6 V·s/cm². Automatic calibration of ion mobility was set ON.

For analyzing PTMs, the Astral was operated in data-dependent mode, performing a full scan in the Orbitrap every 0.7 s (m/z range 375–1,500 m/z, resolution 240,000, AGC target 1,000,000, maximum injection time 10 ms) followed by MS/MS scans of the most abundant ions. MS/MS spectra were acquired in the Astral analyzer using HCD collision energy of 30, isolation width of 1.6 m/z, minimum intensity of 5,000 and maximum injection time of 10 ms. Fragment ions ranging from 80–1,600 m/z were acquired. Precursor ions selected for fragmentation (include charge state 2–6) were excluded for 30 s. The monoisotopic precursor selection filter and exclude isotopes feature were enabled.

Some PTMs and in vitro (de)acetylation assays were analyzed on the Orbitrap Exploris 480 mass spectrometer, which was operated in data-dependent mode, performing a full scan (m/z range 350–1,200, resolution 60,000, AGC target 1,000,000) followed by MS/MS scans of the most abundant ions for a cycle time of 1 s. MS/MS spectra were acquired using HCD collision energy of 30, isolation width of 1.4 m/z, orbitrap resolution of 30,000, AGC target 200,000, minimum intensity of 50,000 and maximum injection time of 80 ms. Precursor ions selected for fragmentation (include charge state 2–6) were excluded for 10 s. The monoisotopic precursor selection filter and exclude isotopes feature were enabled.

### Data Processing protocol

Astral and TimsTOF DIA data were analyzed in Spectronaut 20.3. [<u>25724911</u>] (Biognosys). Trypsin/P was specified as a proteolytic enzyme and up to 2 missed cleavages were allowed in the Pulsar directDIA+ search. Dynamic mass tolerance was applied for calibration and main search. The search was performed against organism-specific databases as described in PRIDE data repository supplemented with common contaminants. Beta-methylthiolation of cysteine was searched as fixed modification, whereas oxidation of methionine and acetylation at protein N-termini were defined as variable modifications. Peptides with a length between 7 and 52 amino acids were considered and results were filtered using Spectronaut default filtering criteria (Precursor Qvalue <0.01, Precursor PEP <0.2, Protein Qvalue <0.01 per Experiment and <0.05 per Run, Protein PEP <0.75). Quantification was performed as specified in Biognosys BGS Factory Default settings, grouping peptides by stripped sequence and performing protein inference using IDPicker.

Spectronaut results were exported using Pivot reports on the protein and peptide level and converted to Microsoft Excel files using VBC in-house software MS2Go [https://ms.imp.ac.at/?action=ms2go]. For DIA data MS2Go utilizes the python library msReport (developed at the Max Perutz Labs Proteomics Facility) for data processing. Missing values were imputed with values obtained from a log-normal distribution with a mean of 100. To compensate for different protein lengths, protein quantification was then normalized using iBAQ [<u>21593866</u>]. For organism-level protein abundance analysis, only proteins with at least one quantified peptide per sample were retained and proteins derived from symbionts or potential contaminants not assigned to the target organism were excluded.

To compare histone-associated protein abundances across different species, we collected proteome datasets from public repositories that cover a wide range of archaeal and eukaryotic lineages (Supplementary Table 8). Publicly available iBAQ proteome data were retrieved from PaxDb v6 [<u>41182819</u>]. For datasets lacking pre-computed iBAQ values, raw mass spectrometry files were re-analyzed using MaxQuant v2.7.3.0 [<u>27809316</u>]. Parameter files were generated using the command-line interface (‘MaxQuantCmd.dll’) with standard DDA settings (LCMSType: ST; instrument type: Orbitrap). iBAQ quantification was enabled by modifying the generated parameter file (‘mqpar.xml’) prior to execution. Searches were performed against organism-specific FASTA databases formatted with the ‘>prefix|accession|description’ header convention. Analyses were run on the CLIP HPC environment.

For analysis of PTMs the RAW-files were loaded into Proteome Discoverer (version 3.2.0.450, Thermo Scientific). All MS/MS spectra were searched using MSAmanda version 3.2.22.93 (<u>24909410</u>). Peptide and fragment mass tolerances were set to ±10 ppm with up to 2 missed cleavages, using Trypsin, Arg-C, Glu-C, or Chymotrypsin as enzyme specificity without proline restriction. For propionylated peptides, either Trypsin with 5 missed cleavages or Arg-C with 2 missed cleavages and enzyme semispecificity (cleavage only on one side, specific after R) was used. Searches were performed against organism-specific databases (Supplementary Table 9) supplemented with common contaminants. Carbamidomethylation of cysteine was set as a fixed modification for alkylated samples. Variable modifications included: oxidation of methionine, mono- and dimethylation on arginine and lysine (in some cases also on aspartic acid, glutamic acid, and the protein N-terminus), trimethylation on lysine, acetylation on lysines and the protein N-terminus, propionylation and propionylation plus methylation on lysines and the protein N-terminus (and for doubly propionylated samples on all N-termini), formylation on lysines (and in some cases on peptide N-termini), and glutamine-to-pyroglutamate conversion at peptide N-terminal glutamine. In selected searches, phosphorylation on serine, threonine, and tyrosine and ubiquitination on lysines were also included. PTM site localization was performed using ptmRS, based on phosphoRS^70^. The result was filtered to 1 % FDR on PSM and protein level using the Percolator algorithm^71^ integrated in Proteome Discoverer. Additionally, an Amanda score cut-off of at least 150 was applied. Proteins were filtered to be identified by a minimum of 2 PSMs in at least 1 sample. Protein areas were computed in IMP-apQuant^72^ by summing up unique and razor peptides. Resulting protein areas were normalized using intensity-based absolute quantification (iBAQ)^73^. Match-between-runs (MBR) was applied for peptides with high confident peak area that were identified by MS/MS spectra in at least one run. Proteins were filtered to be identified by a minimum of 3 quantified peptides. Spectra of modified peptides were validated manually.

For the identification and relative quantification of synthetic peptides in (de)acetylation reactions, DDA data were processed, quantified on MS1 level and manually evaluated using Skyline (64-bit, v22.2.0.351).

### ‘KowaHeim1’ environmental sample

An iron-rich brown biofilm sample was collected from the production well of the Kowakubi hot spring (39.5584° N, 140.2823° E) in 2023. The sample was immediately frozen on dry ice at the sampling site and stored at −80 °C until further processing. DNA and RNA were co-extracted using the ZymoBIOMICS DNA/RNA Miniprep Kit (Zymo Research, CA, USA) according to the manufacturer’s instructions. A metagenomic sequencing library was prepared using the Illumina DNA Prep (M) Tagmentation kit with Nextera DNA CD Indexes (Illumina, CA, USA) and sequenced (2 × 150 bp paired-end) on the Illumina NovaSeq X platform at Macrogen Japan. Metagenome-assembled genomes (MAGs) were reconstructed from the production well biofilm metagenome as described previously^74^. A MAG affiliated with the Heimdallarchaeota was further refined using the CLC Genome Finishing Module as described elsewhere^75^. For metaproteomic analysis of the KowaHeim1 in the well biofilm, total proteins were extracted from the frozen well biofilm sample as described above.

Taxonomic assignment of the archaeal metagenome-assembled genome (MAG) was performed using GTDB-Tk v2 with the classify workflow (‘classify_wf’) against the Genome Taxonomy Database (GTDB) release 232 (r232) reference data^76^. GTDB-Tk identifies, aligns and places genomes into the GTDB reference tree using a combination of marker-gene identification, multiple sequence alignment and placement with pplacer. The resulting classification assigned the MAG (WAKA01) to the family Kariarchaeaceae within the Asgard archaea. The placement tree was visualized using PearTree (https://peartree.live/).

### Transcriptome analysis

Publicly available transcriptome datasets for *Ca.* Lokiarchaeum ossiferum B35 (Loki-B35) and *Ca.* Margulisarchaeum peptidiphilum HC1 (Hod-HC1) was obtained from the NCBI Sequence Read Archive (Loki-B35 SRA accessions: SRX27494461-SRX27494466; Hod-HC1 SRA accession: DRR620298)^28,29^. Raw paired-end reads were quality-filtered using Trimmomatic v0.39^77^ with adapter trimming (TruSeq3-PE-2 adapters; mismatch threshold 2, palindrome clip threshold 30, simple clip threshold 10) and quality filtering parameters: ‘LEADING:3, TRAILING:3, SLIDINGWINDOW:4:20, MINLEN:36’. Transcript abundance was quantified using Salmon v1.10.1^78^ in selective-alignment mode with the ‘--validateMappings’ flag against the respective genomes (Loki-B35 GenBank accession: CP104013.1; Hod-HC1 GenBank accession: AP029002.1) ^28,29^. Transcript-level abundances were reported as transcripts per million (TPM).

### Immunofluorescence of Loki cells and antibody validation

Peptide synthesized from the N-terminal of tailed histone of the HLoB.1 Loki-B35 histones (C-RKRVTAADVIAAE-OH) was conjugated to keyhole limpet hemocyanin (KLH) using the peptide synthesis service from Vienna Biocenter Core Facilities. Rabbits were immunized against the synthetic peptide by Eurogentec, and then antibodies were purified from serum in-house. To validate the Loki tailed histone antibody, the *E. coli* strain BL21 was transformed with the IPTG-inducible vector, pET-15b, containing each of the HLoA and HLoB Loki histones. Single colonies were grown in 5 mL LB + ampicillin (100 μg/mL) at 37°C overnight. The following day, 300 μL of culture was added to a fresh volume of 5 mL LB + amp and grown for 2 hr, or until reaching an OD_600_ of 0.6. Cultures were induced with IPTG (F.C. 500 μM). After 4 hr, 500 μL of culture was spun down, then boiled in 100 μL of Laemmli buffer for 5 min. Then, 15 μL of sample expressing each type A and B Loki histone was run on 15% acrylamide gels to check the cross-reactivity of the HLoB antibody with HLoA and HLoB Loki histones. The membrane was blocked overnight at 4°C in 5% skim milk powder dissolved in TBS-T. The following day, the purified histone antibody was diluted to 1:1000 in blocking solution and incubated with the membrane for 1 hr on a shaker at RT. The secondary incubation proceeded for 45 min using goat anti-rabbit IgG coupled to HRP at a 1:10,000 dilution.

For immunofluorescence experiments, Loki-B35 cells were fixed with Paraformaldehyde (F.C. 3.7%) for 15 min. Next, the enrichment culture was centrifuged for 5 min at 10,000 × g, the supernatant removed and the pellet resuspended in PBS (1/5^th^ of the original volume). Subsequently the cells were transferred onto poly-L-lysine (0.01%)-coated coverslips and incubated for 60 min. The cells were permeabilized with 0.05% Triton X-100 for 10 min at room temperature and blocked with a 2% BSA solution in PBS-T for 60 min at room temperature. The primary antibody against the HLoB tailed histones (described above) was diluted 1:500 and the staining solution containing the secondary antibody (goat anti-rabbit IgG Alexa Fluor 568, Abcam ab175471) and NucBlue were diluted 1:1000 in PBS-T + 0.1% BSA. Incubation times were either 1 hr (primary) or 0.5 hr (secondary) at room temperature, protected from light with gentle shaking. In between the steps, the coverslips were washed gently with PBS or PBST by changing the solution twice. Finally, the coverslips were mounted on microscopy slides using the VECTASHIELD antifading agent. Fixed cells were imaged with a Nikon Eclipse upright microscope equipped with a 100× Nikon 1.45 PlanApo oil objective. Images were acquired using an LED lamp and dichroic mirrors used for visualization of DAPI (405 nm) and Alexa 568 (568 nm). The exposure time was 20 ms for DAPI and 30 ms for Alexa 568.

### Image analysis

Image analysis was performed with Fiji^79^. For visualization purposes, the contrast of the images was enhanced for best visualization, while data analysis was performed on raw images. To quantify intensities of the secondary antibody signal, we selected cells using phase contrast and DNA channels and then quantified line profiles (line width=5) drawn across the cell body in the 568 nm channel. The reported gray value intensities in boxplots were the maximum intensities measured after background subtraction. To measure intensities along the protrusions of Loki cells, a line was drawn along the protrusion in the phase contrast channel, using the segmented line tool. Next, the line was used to quantify intensity traces in the 405 nm (DNA) and 560 nm (secondary antibody) channels. The measured intensities were then normalized to 0–1 using min–max normalization. Finally, the data was plotted using Jupyter Notebook.

### Clustering-based protein identification and phylogenetic analysis

To reconstruct the evolutionary history of Loki proteins, we constructed orthologous gene clusters (i.e., orthogroups) in 255 representative proteome datasets from a tree of life comprising 18 Bacteria, 217 Archaea (63 Asgard archaea, 52 TACK, 53 Euryarchaeota, 48 DPANN, and *Ca.* Sukunaarchaeum mirabile M16-5), and 20 eukaryotes (Supplementary Table 4). Orthofinder v2.5.2^80^ was used to cluster genes in a non-biased way by comparing each gene to the entire proteome dataset (*n* = 845,790). After testing different parameters, we chose DIAMOND^81^ for a homology search with ‘-S diamond_ultra_sens’ and adjusted five different inflation parameters (‘-I’: 1.5, 1.25, 1.2, 1.15, and 1.05). MAFFT v7.310^82^ was used to align protein sequences based on orthogroups. Individual maximum likelihood gene trees were built with IQ-TREE v2.1.2^83^, which used model selection (‘-m MFP’) and an ultrafast bootstrap approximation approach (1,000 replicates). Protein domains were identified using HMMER v3.4^84^ using the Pfam 37.0 database^85^. The absence and presence of proteins (or domains) after manual correction of proteins were visualized next to the species tree using iToL v6.7^86^.

### Histone sequence similarity network construction

Histone and histone-like protein sequences were identified from orthologous groups (OGs) and by HMM-based searches^84^ using profiles constructed from histone-like proteins. All sequences were combined into a single fasta file for downstream analysis. Pairwise sequence similarity was assessed using an all-versus-all search with DIAMOND v2.1.8^81^ in ultra-sensitive mode. A DIAMOND database was constructed from the merged sequence set, and searches were performed with an E-value threshold of 1e-05. To increase network connectivity, a second search was conducted using a relaxed threshold (E-value 1e-03).

Similarity scores were converted into a graph representation using the Markov Cluster Algorithm (MCL) pipeline^87^. BLAST tabular output (format 6) was transformed into a similarity matrix using ‘mcxload’ with mirrored edges, and clustering was performed with MCL v14.137 using multiple inflation parameters (in a range of 1 to 1.5) to explore clustering granularity. Cluster assignments were extracted using ‘mcxdump’ and processed with a custom Python script to generate Cytoscape-compatible files. Self-hits were removed and pairwise similarities were retained as weighted edges annotated with sequence identity, bit score, and E-value. Node attributes were defined by assigning each protein to its corresponding MCL cluster. Networks were visualized in Cytoscape v3.10.2^88^. Nodes were colored according to cluster membership using discrete mapping, and layouts were generated using force-directed or organic algorithms. The scaling of edge thickness was optional and depended on bit score, while weak connections were filtered based on E-value thresholds.

### Structural-based functional annotation of proteins

To complement sequence-based functional inference, protein structural homology was used to identify distant homologs and refine the functional landscape of Asgard archaeal proteomes. High-confidence three-dimensional protein structures were predicted for representative Asgard archaeal proteomes using AlphaFold2^89^.

Structural similarity searches were performed using Foldseek v2024^90^, which enables rapid and sensitive alignment of large structure sets by encoding tertiary interactions into a structural alphabet. Functional annotations were obtained using two complementary bidirectional approaches: (1) querying predicted Asgard archaeal structures against public repositories to identify characterized homologs, and (2) using experimentally validated or predicted eukaryotic structures as queries to identify structural counterparts within the Asgard archaeal proteome.

To ensure functional reliability and minimize noise from uncharacterized proteins or gene modeling artifacts, predicted structures were systematically compared against the RCSB Protein Data Bank (PDB)^91^]. Only structural alignments to experimentally validated and characterized proteins within the PDB were utilized for high-confidence functional refinement. Cross-referencing between predicted structural models and the PDB facilitated the detection of distant evolutionary relationships that were otherwise inaccessible to traditional sequence-based methods.

### In silico protein-protein interaction comparison using AlphaFold screening

To evaluate histone-associated protein-protein interactions, we performed high-throughput in silico interactome screening using AlphaFold2-Multimer^89,92^ implemented by ColabFold^93^. Predictions were performed in the CLIP HPC environment (https://www.clip.science/) using the high-throughput ColabFold pipeline ‘ht-colabfold’ (https://gitlab.com/BrenneckeLab/ht-colabfold)^94^.

Our primary analysis focused on the *Ca.* Lokiarchaeum ossiferum B35 proteome (*n* = 5,119). To maximize sensitivity for detecting histone regulators and to probe distinct interaction modes, we employed multiple bait configurations, including full-length histones, truncated histone tails, and histone dimers (both homo- and heterodimers). These baits were systematically screened against candidate interaction partners. Candidate interactions were prioritized using a custom interaction score (‘PEAKscore’) defined as the scaled (0-1) inverse of the minimum predicted aligned error (PAE) between chains (see details https://gitlab.com/BrenneckeLab/ht-colabfold). This metric captures the confidence of inter-chain orientation while excluding intrachain PAE, thereby specifically emphasizing interface quality. We applied a stringent threshold of PEAKscore 0.75 to define high-confidence interactions, consistent with benchmarks established from a previous study^94^. To extend the analysis to histone modification contexts, we performed additional targeted screening using AlphaFold3^95^. Each of the three Loki SIR2 paralogs (*Lo*SIR2a, b, and c) was modeled in a complex with HLoB.2 histone bearing individual acetylation marks at lysine residues identified by mass spectrometry. Prediction confidence was evaluated using the interface predicted template modelling score (iPTM) and the predicted template modelling score (pTM) (Supplementary Table 10). Resulting structural models and predicted interfaces were visually inspected and rendered using UCSF ChimeraX^96^.

### Molecular dynamics simulations

AlphaFold3 histone dimer predictions were placed into the HHoB closed hypernucleosome cryo-EM density (EMD-53387) by rigid body fitting in ChimeraX ^32,96^. Six dimers combined with 180 bp of DNA from the same structure (PDB: 9QV6) served as the starting model. All-atom molecular dynamics simulations were carried out using AMBER^97^ with the ff14SB, bsc1, and TIP3P force fields for protein, DNA, and water, respectively. Parameters for modified residues, i.e., mono-methyllysine (MLZ), di-methyllysine (MLY), acetyllysine (ACK), and mono-methylarginine (EME) were retrieved from the wwPDB Chemical Component Dictionary^98^. These were processed with ANTECHAMBER and PARMCHK2 to generate Generalized Amber Force Field (GAFF)-compatible parameters. The resulting parameter and library files were loaded concurrently with standard force field sources during system preparation in TLEAP. Structures were protonated and solvated in cubic TIP3P boxes with a 20 Å buffer around the solute and charge-neutralized using sodium and chloride ions, with additional ions (equivalent to 100 mM NaCl and 10 mM MgCl_2_) added to approximate physiological conditions. Hydrogen mass repartitioning was applied using PARMED.

Energy minimization was performed in two sequential stages: the first (1,000 steps) restrained all non-solvent atoms to allow solvent relaxation, and the second (2,500 steps) allowed full system relaxation without restraints. Minimized structures were then heated from 0 to 300 K over 100 ps under constant volume conditions with positional restraints on the solute using Langevin dynamics. A brief NVT equilibration (200 ps) followed with light solute restraints, succeeded by NPT equilibration (1 ns) at 300 K and 1 atm with the Berendsen barostat (τ_P_ = 2.0 ps) and no restraints. Production simulations were run for 500 ns under NPT conditions at 300 K and 1 atm in 4 fs steps, with SHAKE constraints applied to all bonds involving hydrogen. Complexes were simulated with wild type sequences (WT) and mutants (MUT) incorporating all the detected PTMs (Extended Data Fig. 5). Simulations were performed in triplicate by initializing each replicate with a different random seed during the heating phase.

Trajectory analyses were performed in Python using the MDAnalysis, NumPy, SciPy, and Matplotlib libraries^99–101^. Root-mean-square deviation (RMSD) of the protein backbone Cα atoms was calculated relative to the initial structure for each trajectory after least-squares superposition (translational and rotational fitting) to remove overall rigid-body motion (Supplementary Figure 6). To quantify nucleosome breathing dynamics, inter-strand distances were calculated across the simulation trajectories. Ten symmetrically selected base-pair residue pairs spanning the center of the DNA were utilized as spatial probes, tracking the distance between opposing C2’ atoms. Mean pairwise distances and their corresponding standard deviations were computed at each frame across the three independent replicates. The mean pairwise distance reports the average separation between neighboring superhelical turns at the probed base pairs at each time point in the simulation (Supplementary Figure 6).

To characterize DNA breathing dynamics, the mean C2’–C2’ distance signal was smoothed using a uniform filter and analyzed in two ways: a threshold-crossing analysis (mean + 0.5σ) was performed to quantify the rate of breathing events per nanosecond and the autocorrelation function was computed to extract the characteristic decorrelation timescale (τ₁/ₑ). The crossing rate captures how often the DNA superhelical turn distance is beyond the average of the simulation from the wild type (Supplementary Figure 6). The ACF decorrelation time describes how long a given breathing fluctuation persists (Supplementary Figure 6). To examine position-dependent dynamics, the ten probe pairs were grouped into two regions: “ends” (pairs 1–3 and 8–10) and “middle” (pairs 4–7). Breathing rates and decorrelation times were averaged within each region and compared between conditions using the Mann-Whitney U test (two-sided).

Free-energy profiles along the C2′–C2′ inter-strand distance were obtained by Boltzmann inversion of the distance distributions, G(d) = –kT ln P(d), with kT = 0.592 kcal/mol (RT at 300 K). For each replicate, distances from all base pairs within a given region ("ends" or "middle") were pooled and binned into a common 400-bin histogram, and the resulting per-replicate profiles were each zeroed to their global minimum before averaging across replicates. The free-energy profile denotes how often a given interstrand distance is sampled over the simulation, with low-energy regions corresponding to conformations occupied most frequently and higher-energy regions corresponding to rarer, more transient states (Supplementary Figure 6).

### Recombinant protein expression and purification for *Lo*SIR2s and *Lo*HATs

Genes encoding candidate chromatin-modifying enzymes were codon-optimized for expression in *Escherichia coli* and synthesized by direct synthesis service from commercial vendors (Twist Bioscience, Genscript). Archaeal and bacterial chromatin-modifying enzymes (SIR2 deacetylases and HAT acetyltransferases) were expressed as N-terminal tag (e.g., 14×His-GST, Myc) fusion proteins with PreScission protease (3C) cleavage sites in *Escherichia coli* BL21 (DE3). Genes were codon-optimized for *E. coli*, synthesized, and cloned into pETM-30 derivative vectors with kanamycin resistance. Expression was induced by autoinduction in Terrific Broth containing 1.5% (w/v) lactose at 25°C for 24 hr. Cells were lysed by sonication, and His-tagged proteins were purified by Ni-Sepharose affinity chromatography using buffers containing 50 mM Tris-HCl pH 8.0, 300 mM NaCl, 0.5 mM TCEP, and 40–500 mM imidazole. Protein concentrations were estimated by SDS-PAGE against BSA standards. Construct details are provided in Supplementary Table 11.

### Fluorophore deacetylation assays

Deacetylase activity of recombinant Loki SIR2 proteins (*Lo*SIR2a–c) was quantified using the SIRT1 Activity Assay Kit (Abcam, ab156065). Deacetylation of the substrate by SIR2 deacetylase renders the peptide susceptible to cleavage by a peptidase (‘developer’), which releases a free fluorophore and generates a fluorescent signal that is proportional to deacetylase activity. Reactions were set up in black 96-well microplates, with a total volume of 50 μL per well, containing assay buffer, fluoro-substrate peptide (component ‘#2’), NAD^+^ cofactor, and developer solution. Reactions were initiated by the addition of 5 μL of purified recombinant protein at five different concentrations. Recombinant human SIRT1 (UniProt: Q96EB6) served as a positive control. Negative controls included reactions without enzyme, without NAD^+^, or with heat-inactivated enzyme. The fluoro-deacetylated peptide (component ‘#3’) was used as a developer control. Plates were incubated at 20°C for 30–60 min with continuous fluorescence readings at excitation/emission wavelengths of 360/460 nm using a Synergy 4 plate reader (BioTek). All reactions were performed in duplicate. Background fluorescence was subtracted from all readings, and deacetylase activity was plotted as a function of protein concentration.

### In vitro acetylation and deacetylation assays coupled to mass spectrometry

Enzymatic activity of candidate histone acetyltransferases (HATs) and sirtuin-like deacetylases was assessed using synthetic histone tail peptides as substrates. Histone tail peptides derived from Loki-B35 and yeast H4 were synthesized in both unmodified and acetylated forms with specific acetylation states at defined lysine residues (Supplementary Table 12). Recombinant HAT candidates were incubated with unmodified histone peptides, whereas the Sir2 homologs (*Lo*SIR2s) were incubated with acetylated histone peptides. Reactions were performed under optimized conditions and terminated prior to analysis. In parallel, reaction products were analyzed by liquid chromatography-tandem mass spectrometry (LC-MS/MS) to confirm site-specific modifications.

We measured the enzymatic activities of candidate sirtuin deacetylases (SIR2) and histone acetyltransferases (HATs) using synthetic histone tail peptides from Loki archaeal histones and yeast histones H4. Peptides were made with specific acetylation states at certain lysine residues and dissolved in 50 mM Tris-HCl (pH 7.5) to make 1-5 mM stock solutions.

For acetylation assays, a master mix with either acetylated or unacetylated peptides (F.C. 100 μM), acetyl-CoA (F.C. 100 μM; Merck 10101893001), and reaction buffer (50 mM Tris-HCl pH 8, 50 mM NaCl, 1 mM DTT—up to the final reaction volume) was warmed to 30°C before starting the first time point. Purified HATs (F.C. 0.5 μM) were then added to the reaction mixture. Enzymes in heat-inactive enzyme controls were heated at 95°C for 5 min before being added. Volumes were removed at the appropriate time point and acid-quenched immediately in a separate Eppendorf tube using trifluoroacetic acid (F.C. 0.1%). Acid-quenched samples were mixed thoroughly, then placed on ice. Samples were submitted to LC-MS in solution. Measurement details are provided in Supplementary Table 13.

For deacetylation assays, acetylated peptides (100 µM) were mixed with purified recombinant SIR2 (0.5 µM) in a reaction buffer that contained 50 mM Tris-HCl (pH 8.0–9.0), 1–4 mM MgCl_2_, 0.2–1 mM DTT, and 1 mM NAD^+^. Reactions were done at 20°C or 37°C in a total volume of 30 µl. Reactions were sampled at 0, 15, 30, 60, and 120 min and terminated by addition of trifluoroacetic acid (0.1% final concentration) followed by immediate cooling on ice. In selected experiments, reactions were alternatively stopped by heat inactivation (95°C, 5 min) and centrifugation. Reaction products were analyzed by liquid chromatography–tandem mass spectrometry (LC-MS/MS). Negative controls included reactions lacking enzyme or lacking a cofactor (NAD^+^ or acetyl-CoA). Yeast histone H4 peptides served as positive control substrates for SIR2 activity.

### CRASH assay for testing the functions of SIR2 homologs

To assess the functional conservation of SIR2 homologues, we employed the CRASH (Cre-reported altered states of heterochromatin) assay^53^ in *Saccharomyces cerevisiae*. In this system, functional silencing of the heterothallic mating type loci (*HMLα/HMRa*) by Sir2 prevents expression of *Cre* integrated there. However, transient losses of silencing trigger an irreversible, Cre-mediated switch from RFP to GFP expression of a kanMX6-containing reporter cassette located within euchromatin. Such irreversible switching is then used to quantify silencing efficiency through flow cytometry and qualitatively observed as sectoring on agar plates.

Chimeric SIR2 constructs were engineered by synthesizing codon-optimized gene fragments containing the catalytic domain of archaeal Sir2 homologs and then fusing these to the *S. cerevisiae* Sir2 targeting domain (DUF592 domain), which directs the protein to *HMLα/HMRa*. This fusion step was performed by Gibson assembly into the pNX3b-Myc13 (Addgene 78367) integration vector for C-terminal Myc-tagging and hygromycin selection. All chimeric archaeal Sir2-integration constructs, along with catalytic-dead negative controls, are detailed in Supplementary Table 14.

Sir2 fusion constructs were integrated at the endogenous *SIR2* locus of the CRASH reporter strain (YZH_1345) by PCR-mediated homologous recombination^102^. Transformants were plated on YPD supplemented with 100 µg/ml hygromycin, and genotyping was performed by colony PCR followed by Sanger sequencing of the *SIR2* locus. Genetically verified colonies were then replicated onto YPD supplemented with 200 µg/ml to assess whether they still maintained the reporter cassette. For those strains for which we did not recover any colonies maintaining the reporter cassette, expression of the archaeal Sir2 homologue was verified by immunoblot using anti-Myc 9E10 (IMP Molecular Biology Service) for detection. All strains used in this study are reported in Supplementary Table 15.

To quantify the apparent silencing frequency of archaeal Sir2 homologues, we measured the fraction of GFP-only, RFP-only, and both GFP- and RFP-expressing cells by flow cytometry^103^. Briefly, cells were pre-grown to saturation in 200 µl YPD supplemented with 200 µg/ml G418, except for strains for which we were unable to recover any silencing-competent colonies, which were pre-cultured in YPD alone. These pre-cultures were then diluted 1:100 in YPD and OD_600_ was monitored until they reached mid-log (OD_600_ ∼ 0.5). Cells were then fixed in 4% paraformaldehyde for 30–45 min, washed once with PBS, and stored in PBS at 4 °C until flow cytometry measurement. Flow cytometry was performed on an IntelliCyt iQue Screener Plus at the BioOptics facility, with at least 10,000 events captured.

To qualitatively assess the silencing frequency of Sir2 homologues, we further detected colony sectoring by fluorescence microscopy. Briefly, saturated pre-cultured CRASH strains grown in YPD with or without 200 µg/ml G418 were diluted 1:20,000 in sterile water, and then 300 µl was plated onto 9 cm petri plates containing 15 ml of SD-Trp growth medium with 1% agar. Thin, low-agar, and tryptophan-dropout mediums were used to reduce background fluorescence. Imaging was performed on a Leica AxioZoom V16 Stereomicroscope with sCMOS camera at the BioOptics facility.

## Data availability

Mass spectrometry proteomics data have been deposited in the PRIDE repository: whole-cell lysate proteomes, histone-enriched proteomes, and (de)acetylation mass spectrometry assay. The draft genome of *Candidatus* Heimdallarchaeota archaeon ‘KowaHeim1’ has been deposited at DDBJ/ENA/GenBank under the accession number JBZXTT000000000. Source data for all main and supplementary figures are available at the Dryad dataset repository (https://doi.org/10.5061/dryad.gqnk98t3w).

## Code availability

The codes used in this study are available in the Dryad repository at https://doi.org/10.5061/dryad.gqnk98t3w. The high-throughput ColabFold pipeline used for protein-protein interaction screening is available at https://gitlab.com/BrenneckeLab/ht-colabfold.

## Acknowledgments

We thank the Vienna BioCenter Core Facilities, in particular the Proteomics Facility and the Protein Technologies Facility, for their expertise, helpful discussions, and recombinant protein production, and the Molecular Biology Service, Media Kitchen, and Peptide Synthesis Service for reagents, cloning, sequencing, and peptide synthesis. We thank the BioOptics facility of the IMP/IMBA/GMI for microscopy and flow cytometry support. We thank the members of the Berger group for their support throughout this project, especially Zdravko J. Lorković for antibody and peptide preparation and Nenad Grujic for harvesting Asgard archaeal cultures. We thank Gabriel Thornes and Masaru Sanari from the Schleper group for helping with immunostaining, and Fatma Baraket, Maximilian Dreer, Thomas Pribasnig, and Andrea Maltis for contributions to archaeal cultures and sample preparation. We thank Jian Yi Kok (*S. pombe*), Kensuke Kataoka (*T. thermophila*), Tatsuo Kanno and Amellie Gonon (*A. thaliana*), Miyuki Ogawara, Yumi Saito, and Masami Isozaki (*Ca.* M. peptidiphilum HC1), Pamela Vetrano and Silvia Ramundo (*C. reinhardtii*) for providing cells and biomass. We thank Christoph Büschl for assisting with AlphaFold-Multimer predictions, Wen-Cong Huang for advice on archaeal taxon sampling, and Takeshi Kakegawa and the owner of Kowakubi Onsen Syohoen for the environmental sample of ‘KowaHeim1’. We thank Carol Featherstone from Life Science Editors for editorial assistance. We thank Tobias Viehböck and the members of the EvoChromo consortium (https://evochromo.org/) for insightful discussions, and Yoav Voichek, Pierre Bourguet, Arie Fridrich, and Sean A. Montgomery for comments on the manuscript.

## Contributions

F.B., F.K.M.S. and C.S. acquired funding and supervised the project. F.B. and C.H.C. conceptualized the study and coordinated the work. K.H., N.P., N.M., L.H.H., H.I., S.I., J.N.H., A.S. and I.A.Z. provided archaeal biomass for proteomics. C.S. supervised most of the archaeal cultivation. H.I. and S.I. provided ‘KowaHeim1’ genomic data. C.H.C. and C.S.R. processed proteomics samples and performed in vitro biochemical assays and hPTM experiments. Z.H.H. designed yeast experiments. Z.H.H. and J.C. performed CRASH assays and microscopic imaging of complementation. P.R. performed immunofluorescence microscopy with Loki-B35 tailed histone antibody. R.I., G.K. and E.R. performed mass spectrometry and hPTM analysis. C.H.C., C.S.R., M.H., and F.K.M.S. performed AlphaFold predictions and structural analysis. H.M.R. and S.O.D. performed molecular dynamics simulations. C.H.C., C.S.R. and N.A.T.I. performed phylogenetic and bioinformatic analyses. C.H.C., C.S.R., Z.H.H, and F.B. prepared figures and wrote the manuscript. All authors reviewed, edited and approved the manuscript.

## Supplementary Figures

**Supplementary Figure 1. Overview of histone distribution and taxonomic sampling across 255 taxa in the tree of life.** Taxa are organized along an approximate species tree inferred using OrthoFinder’s STAG/STRIDE pipeline from gene tree reconciliation. Note that this tree is used solely to organize taxa by major clade for visualization of histone distribution patterns; inter-clade branching order does not reflect current phylogenomic consensus on deep phylogenetic relationships. Red arrowheads indicate species for which whole-cell proteome data were generated in this study. Filled squares denote the presence of nucleosomal histones, while open squares denote their absence. Taxonomic abbreviations for all species names are provided in Supplementary Table 4.

**Supplementary Figure 2. Phylogenetic placement of the metagenome-assembled genome ‘KowaHeim1’.** The reference tree was inferred from the GTDB release 232 (r232) archaeal backbone using GTDB-Tk. The tree shows Heimdallarchaeia with four major lineages: Hodarchaeales, Gerdarchaeales, Kariarchaeaceae and Heimdallarchaeaceae. The MAG recovered in this study was classified within the family Kariarchaeaceae and is labeled *‘*KowaHeim1’ (red text, blue box). Leaf labels correspond to GTDB genome accession numbers. The double slash (//) at the root denotes a truncated branch.

**Supplementary Figure 3. Comparative analysis of histone tail properties in Asgard archaea and eukaryotes.** Scatter plots comparing N-terminal tail characteristics across Asgard archaeal histone families and eukaryotic core histones (H2A, H2B, H3, H4). Top: Tail length (amino acids, log_2_ scale). Middle: Tail lysine content (% Lys). Bottom: Isoelectric point (pI). Orange dots represent individual histone sequences; red horizontal lines indicate median values. Sample sizes (n) are shown above each group. Asgard histone tails show considerable diversity: HMpB/HLoB-type histones exhibit long, lysine-rich tails (>30% Lys, ∼16–32 aa) with high pI (∼10.5), resembling eukaryotic histones. In contrast, canonical archaeal histones (e.g., HMpA/C, HLoA) have shorter tails with lower lysine content. Eukaryotic histones display longer tails (32-64 aa) with moderate lysine content (15-32%) and high pI (11–11.5). Statistical comparisons were performed using two-sided Mann-Whitney U tests with Benjamini-Hochberg false discovery rate (FDR) correction for multiple testing. Significance levels: ‘**’ (p<0.01), ‘*’ (p<0.05), ‘n.s.’ (not significant; p>=0.05).

**Supplementary Figure 4. Antibody validation of HLoB histone variants.** Simplified phylogenetic tree showing the relationship between *Ca.* Lokiarchaeum ossiferum B35 histone variants, including HLoA (short-tailed, red clade) and HLoB (long-tailed, blue clade). Western blot (bottom) using an anti-HLoB.1 antibody raised against the N-terminal tail region (C-RKRVTAADVIAAE-OH) demonstrates specific recognition of HLoB recombinant proteins (HLoB.1, HLoB.2, HLoB.3; marked with asterisks) but not HLoA recombinant proteins (HLoA.1, HLoA.2, HLoA.3a, HLoA.4a/b).

**Supplementary Figure 5. Colocalization of DNA and histone signals in Asgard archaeal cells.** Immunofluorescence microscopy showing spatial distribution of DNA (cyan, NucBlue staining) and tailed histones (yellow, detected with anti-HLoB antibody) in *Ca.* Lokiarchaeum ossiferum B35 cells. Left: Phase contrast (PC) image showing cell morphology with characteristic protrusions. Middle: Fluorescence channels showing DNA (cyan) and histone antibody signals (yellow) with overlapping distribution in cell bodies and protrusions. Orange lines indicate representative intensity measurement paths from the cell body into the protrusions. Right: Quantification of normalized fluorescence intensity along measurement paths shows that DNA and histone signals co-decrease from the cell body (position 0 µm) toward protrusion tips, consistent with chromatin localization throughout cell bodies. Data represent mean ± s.d. Scale bar, 5 µm.

**Supplementary Figure 6. Effects of detected post-translation modifications (PTMs) on hypernucleosome structures in molecular dynamics simulations.** Histones HTkA from *T. kodakarensis*, HMfB from *M. fervidus*, HLoA.3a, and HLoB.2 from Loki-B35 were studied. All analyses compare complexes with wild-type protein (WT, black-grey) and mutants with all detected PTMs incorporated (MUT, red-pink). (a) RMSD curves. (b) Mean pairwise distances of 10 DNA pairs. (c) Snapshots of the last frame from three runs. (d) Free energy curves at the end and middle regions. (e) Crossing rate from the mean distance of WT and decorrelation time calculated from the autocorrelation function. All data is presented as a mean ± s.e.m. (*n* = 3).

**Supplementary Figure 7. Additional MS/MS validation of histone tail post-translational modifications in Asgard archaea. a**, MS/MS spectrum and fragment ion table for the Loki-B36 HLoB tailed histone peptide KKAAPKKAAKPATR showing possible acetylation at K1 (HLoB_[10-23]_K9ac). Propionylation (pr) of unmodified lysine residues (K2, K6, K7, K10) results from chemical derivatization during sample preparation. **b**, MS/MS spectrum and fragment ion table for the Hod-HC1 HMpB tailed histone peptide KATKPAPK showing methylation at K4 (HMpB.1_[6-13]_K8me1). The diagnostic b_4_^+^ and y_5_^+^ ion pair localizes the modification to K4. In both panels, fragment ion abundance is plotted at various mass-to-charge ratios (*m/z*). The b and y ions and their losses of H_2_O are marked in red and blue, respectively; precursor ions in green; unassigned peaks in light grey. Nonessential ion labels have been omitted for readability. Fragment ion tables list theoretical *m/z* values for all detected ion series. ac, acetylation; pr, propionylation; me, methylation. Histone residues are numbered canonically from the second residue, as the initiator methionine is absent in the mature polypeptide.

**Supplementary Figure 8. AlphaFold structural prediction of Asgard histone writer. a-d**, AlphaFold2 (AF2) predictions of *Lo*HAT2–HLoB.2 complexes. **a**, Superimposed AF2 models of two most distinct predictions (rank_2 and rank_5) of *Lo*HAT2 (two shades of pink) engaging the N-terminal tail of HLoB.2 (two shades of green) displays high prediction confidence in the protein core fold of both LoHAT2 and HLoB.2 together with their interaction interface. Relative position of histone is variable, due to highly flexible histone tail. As displayed also in Figure 3c, conserved catalytic interface residues on *Lo*HAT2 (orange: Gly106–Ser111 (GKGLMS), Glu95, Asn138) related to acetyl-CoA binding sites are positioned to contact the first accessible lysine on the histone tail (red: Lys3). Models shown in two orientations. **b**, Highest rank model from a, (rank_2) colored by the Predicted Local Distance Difference Test (pLDDT) score from AF2 and shown in two orientations. **c**, Predicted Aligned Error (PAE) plots from AF2 showing the *Lo*HAT2–HLoB.2 interaction prediction. **d**, Highest rank model from a, (rank_2) of *Lo*HAT2 and truncated HLoB.2 model showing the histone tail interacting residue (Lys3), displayed as atom, together with flanking residues. Cartoon representation (left) and surface representation of *Lo*HAT2 (right) visualizing the predicted catalytic cleft, where histone tail is binding. **e,** Experimentally determined structure of human N-terminal acetyltransferase type D (NatD) (light blue) and part of H4 tail (dark green) (PDB: 4U9W) showing the histone tail interacting residue (Ser1) displayed as atom. Cartoon representation (left) and surface representation of NatD visualize the predicted catalytic cleft, where the histone tail is binding, similarly as observed on the predicted *Lo*HAT2 model.

**Supplementary Figure 9. In vitro HAT activity assays using mass spectrometry**. In vitro HAT activity assays using synthetic peptides from the Loki histone tail (HLoB.2_[2-27_) or the yeast histone H4 tail (HHF1_[2-30]_) as substrates were incubated with Loki HAT enzymes (*Lo*HAT1 and *Lo*HAT2). Mass spectrometry reveals that *Lo*HAT2 exhibits robust acetyltransferase activity, generating approximately a 10-fold higher acetylated peptide signal after 120 minutes compared to unincubated controls.

**Supplementary Figure 10. Multiple sequence alignment and conservation of SIR2 family deacetylases.** Top: Multiple sequence alignment of representative SIR2/sirtuin family NAD^+^-dependent deacetylases from eukaryotes, Asgard archaea, TACK, Euryarchaeota, and Bacteria (*n* = 19). Two alignment blocks are shown, spanning the catalytic core. NAD^+^-binding residues (red boxes), proton acceptor residue (purple triangle), and Zn^2+^-coordinating cysteines (blue boxes) are annotated. Red asterisks mark residues mutated to generate the catalytically inactive variant (*Lo*SIR2a-N106A/H123A). Consensus sequence logos above each block indicate positional conservation. Bottom panel: Residue-level conservation index mapped onto yeast Sir2 (UniProt: P06700). The blue line shows the conservation score (inverse Shannon entropy) across the 562 amino acid sequence. The grey line indicates alignment occupancy. Domain architecture is shown above (targeting domain DUF592, olive; SIR2 main domain, blue). Key catalytic residues N345 and H364 are labeled (red asterisks). The high conservation of catalytic and cofactor-binding residues across all domains of life supports an ancestral NAD^+^-dependent deacetylase activity.

**Supplementary Fig. 11. In vitro SIR2 deacetylation assays quantified by mass spectrometry. a,** Deacetylation of fully acetylated Loki histone tail peptide HLoB.2_[2–27]_ (all Kac). The abundance of acetylated peptides (all (9×), 8×, 7×, 6× acetyl) is shown for each enzyme at 15 and 120 min. *Lo*SIR2b progressively converts the fully acetylated substrate into lower acetylation states, whereas *Lo*SIR2a,c and yeast Sir2 show minimal activity on this substrate. **b,** Deacetylation of mono-acetylated Loki histone tail peptide HLoB.2_[2–27]_K9ac. The abundance of mono-acetylated (1× acetyl) and unmodified (no acetyl) peptide is shown. *Lo*SIR2b efficiently removes the single acetylation with the unmodified peptides accumulating by 120 min. **c,** Deacetylation of yeast histone HHF1 tail peptide H4_[2–30]_K16ac. *Lo*SIR2b deacetylates the yeast substrate, whereas other SIR2s show little or no detectable activity under these conditions. In all panels, peptide abundance (y-axis) was measured by mass spectrometry. Coloured backgrounds group enzymes by origin: *Lo*SIR2 paralogs (red, orange, yellow) and yeast Sir2 (blue).

**Supplementary Figure 12. Enzymatic activity of recombinant Asgard archaeal SIR2 deacetylases.** Fluorescence-based deacetylase activity assays were performed for recombinant SIR2 enzymes from Asgard archaea (*Lo*SIR2a, *Lo*SIR2b, *Lo*SIR2c) and human SIRT1 (positive control). Fluorescence (arbitrary units, a.u.) is plotted over time (0–60 min) for varying enzyme concentrations (1× = 4 µg; 1/2×, 1/4×, 1/8×, 1/16×). The 1× concentration corresponds to ∼1.4 µM for *Lo*SIR2a–c (His6-GST fusions, MW ∼56 kDa) and ∼1.0 µM for human SIRT1 (UniProt Q96EB6, MW ∼83 kDa). All measurements were performed in duplicates, and data are presented as mean values. *Lo*SIR2a and *Lo*SIR2b display robust, dose-dependent deacetylase activity comparable to human SIRT1, with fluorescence increasing over time and plateauing at higher enzyme concentrations. *Lo*SIR2c shows minimal activity under standard assay conditions but produces a signal in the deacetylated peptide positive control (‘#3’, dashed grey line), suggesting that the lack of activity may be due to problems with recombinant protein folding or stability rather than an inherent absence of catalytic function. Negative controls (no NAD^+^ and no enzyme blank) show no fluorescence increase, confirming that the observed activity is NAD^+^-dependent.

## Supplementary Tables

**Supplementary Table 1**. Proteome-wide iBAQ abundance and coverage of histone-type and other nucleoid-associated protein families across eukaryotic, archaeal, and bacterial species.

**Supplementary Table 2**. Representative Asgard archaeal histone proteins.

**Supplementary Table 3**. HLoC-type histone sequences identified in this study.

**Supplementary Table 4**. Genomic datasets of 255 representative taxa used in comparative analyses.

**Supplementary Table 5**. Histone-modifying enzymes (writers, erasers) identified in *Ca*. Lokiarchaeum ossiferum B35.

**Supplementary Table 6**. AlphaFold2-based protein–protein interaction clusters of B35 proteins, with functional annotations.

**Supplementary Table 7**. Sample information for cultures used for histone extraction and mass spectrometry in this study.

**Supplementary Table 8**. Whole-cell-lysate (WCL) proteomic datasets and acquisition details across eukaryotic, archaeal, and bacterial species.

**Supplementary Table 9**. Summary of histone post-translational modification (hPTM) proteomic datasets, including peptidase combinations and data sources.

**Supplementary Table 10**. AlphaFold3 prediction results (pTM/ipTM) for *Lo*SIR2 against HLoB.2 histone bearing post-translational modifications.

**Supplementary Table 11**. Recombinant chromatin-modifying enzyme constructs, including plasmid IDs, source organisms, and purification details.

**Supplementary Table 12**. Synthetic peptides used in the in vitro HAT and SIR2 activity assays.

**Supplementary Table 13**. Histone acetyltransferase (HAT) in vitro activity assay data measured by mass spectrometry.

**Supplementary Table 14**. Plasmid constructs used in this study.

**Supplementary Table 15**. *Saccharomyces cerevisiae* strains used in the CRASH complementation assay.

