## Supplementary Figures 1-12 for "Emergence of histone-based chromatin complexity in Asgard archaea"

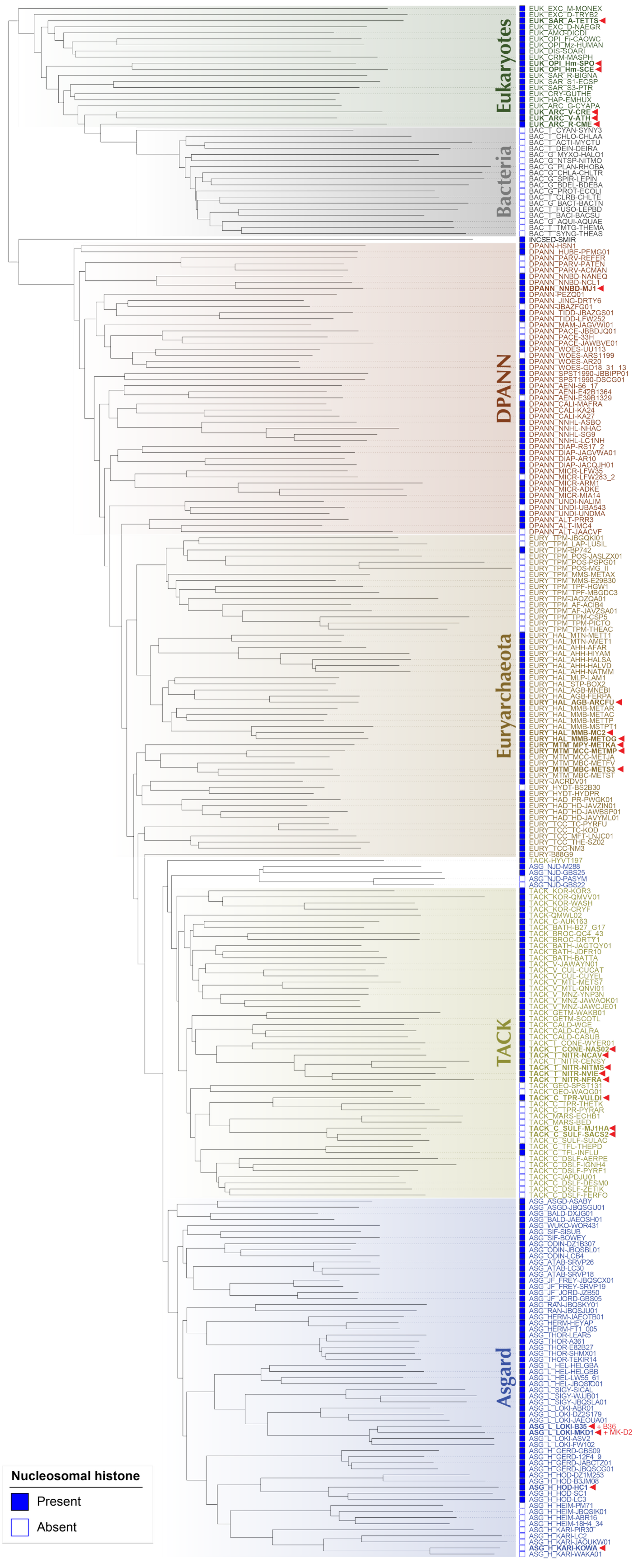
Eukaryotes
Bacteria
DPANN
Euryarchaeota
TACK
Asgard
Nucleosomal histone
Present
Absent

**Supplementary Figure 1. Overview of histone distribution and taxonomic sampling across 255 taxa in the tree of life.** Taxa are organized along an approximate species tree inferred using OrthoFinder's STAG/STRIDE pipeline from gene tree reconciliation. Note that this tree is used solely to organize taxa by major clade for visualization of histone distribution patterns; inter-clade branching order does not reflect current phylogenomic consensus on deep phylogenetic relationships. Red arrowheads indicate species for which whole-cell proteome data were generated in this study. Filled squares denote the presence of nucleosomal histones, while open squares denote their absence. Taxonomic abbreviations for all species names are provided in Supplementary Table 4.

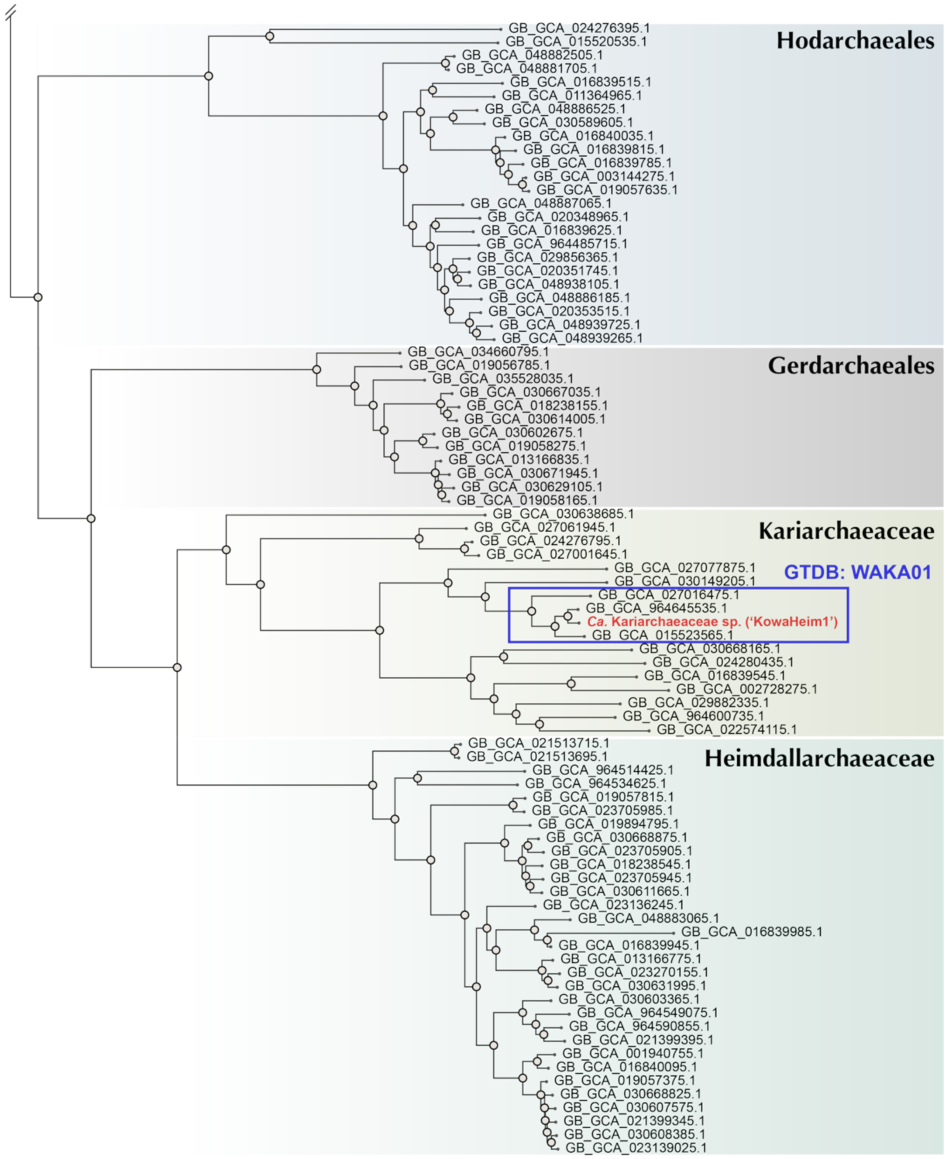


**Supplementary Figure 2. Phylogenetic placement of the metagenome-assembled genome 'KowaHeim1'.** The reference tree was inferred from the GTDB release 232 (r232) archaeal backbone using GTDB-Tk. The tree shows Heimdallarchaeia with four major lineages: Hodarchaeales, Gerdarchaeales, Kariarchaeaceae and Heimdallarchaeaceae. The MAG recovered in this study was classified within the family Kariarchaeaceae and is labeled 'KowaHeim1' (red text, blue box). Leaf labels correspond to GTDB genome accession numbers. The double slash (//) at the root denotes a truncated branch.

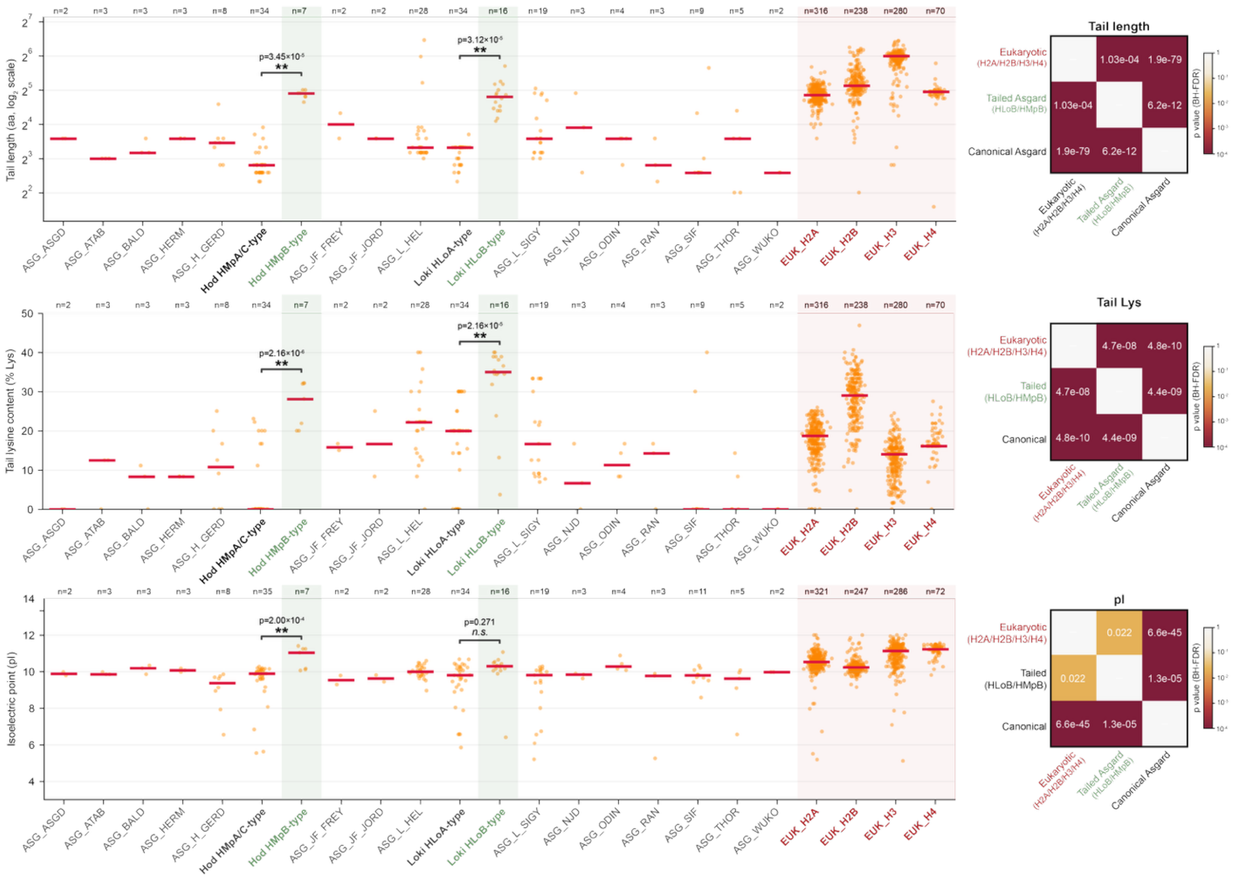


**Supplementary Figure 3. Comparative analysis of histone tail properties in Asgard archaea and eukaryotes.** Scatter plots comparing N-terminal tail characteristics across Asgard archaeal histone families and eukaryotic core histones (H2A, H2B, H3, H4). Top: Tail length (amino acids, $log_2$ scale). Middle: Tail lysine content (% Lys). Bottom: Isoelectric point (pI). Orange dots represent individual histone sequences; red horizontal lines indicate median values. Sample sizes (n) are shown above each group. Asgard histone tails show considerable diversity: HMpB/HLoB-type histones exhibit long, lysine-rich tails (>30% Lys, ~16–32 aa) with high pI (~10.5), resembling eukaryotic histones. In contrast, canonical archaeal histones (e.g., HMpA/C, HLoA) have shorter tails with lower lysine content. Eukaryotic histones display longer tails (32-64 aa) with moderate lysine content (15-32%) and high pI (11–11.5). Statistical comparisons were performed using two-sided Mann-Whitney U tests with Benjamini-Hochberg false discovery rate (FDR) correction for multiple testing. Significance levels: '**' ($p<0.01$), '*' ($p<0.05$), 'n.s.' (not significant; $p>=0.05$).

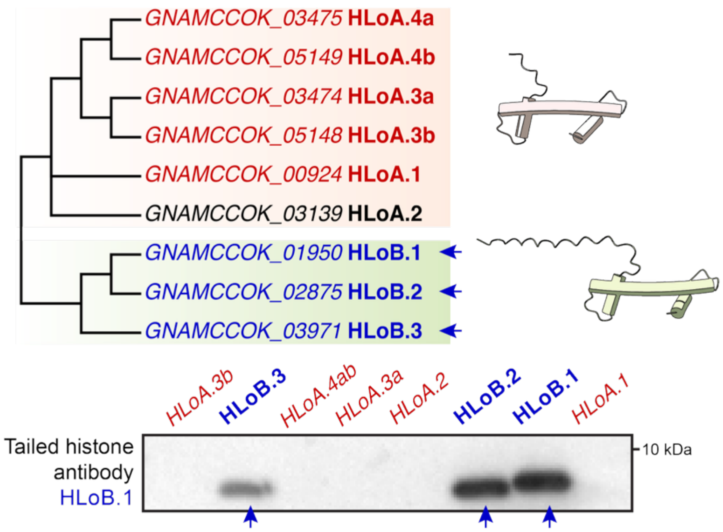


**Supplementary Figure 4. Antibody validation of HLoB histone variants.** Simplified phylogenetic tree showing the relationship between *Ca.* Lokiarchaeum ossiferum B35 histone variants, including HLoA (short-tailed, red clade) and HLoB (long-tailed, blue clade). Western blot (bottom) using an anti-HLoB.1 antibody raised against the N-terminal tail region (C-RKRVTAADVIAAE-OH) demonstrates specific recognition of HLoB recombinant proteins (HLoB.1, HLoB.2, HLoB.3; marked with asterisks) but not HLoA recombinant proteins (HLoA.1, HLoA.2, HLoA.3a, HLoA.4ab).

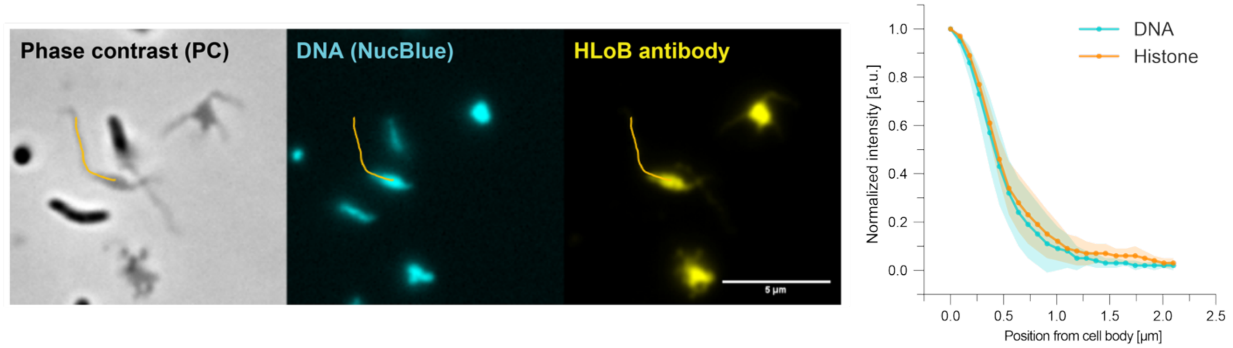


**Supplementary Figure 5. Colocalization of DNA and histone signals in Asgard archaeal cells.** Immunofluorescence microscopy showing spatial distribution of DNA (cyan, NucBlue staining) and tailed histones (yellow, detected with anti-HLoB antibody) in *Ca.* Lokiarchaeum ossiferum B35 cells. Left: Phase contrast (PC) image showing cell morphology with characteristic protrusions. Middle: Fluorescence channels showing DNA (cyan) and histone antibody signals (yellow) with overlapping distribution in cell bodies and protrusions. Orange lines indicate representative intensity measurement paths from the cell body into the protrusions. Right: Quantification of normalized fluorescence intensity along measurement paths shows that DNA and histone signals co-decrease from the cell body (position 0 µm) toward protrusion tips, consistent with chromatin localization throughout cell bodies. Data represent mean ± s.d. Scale bar, 5 µm.

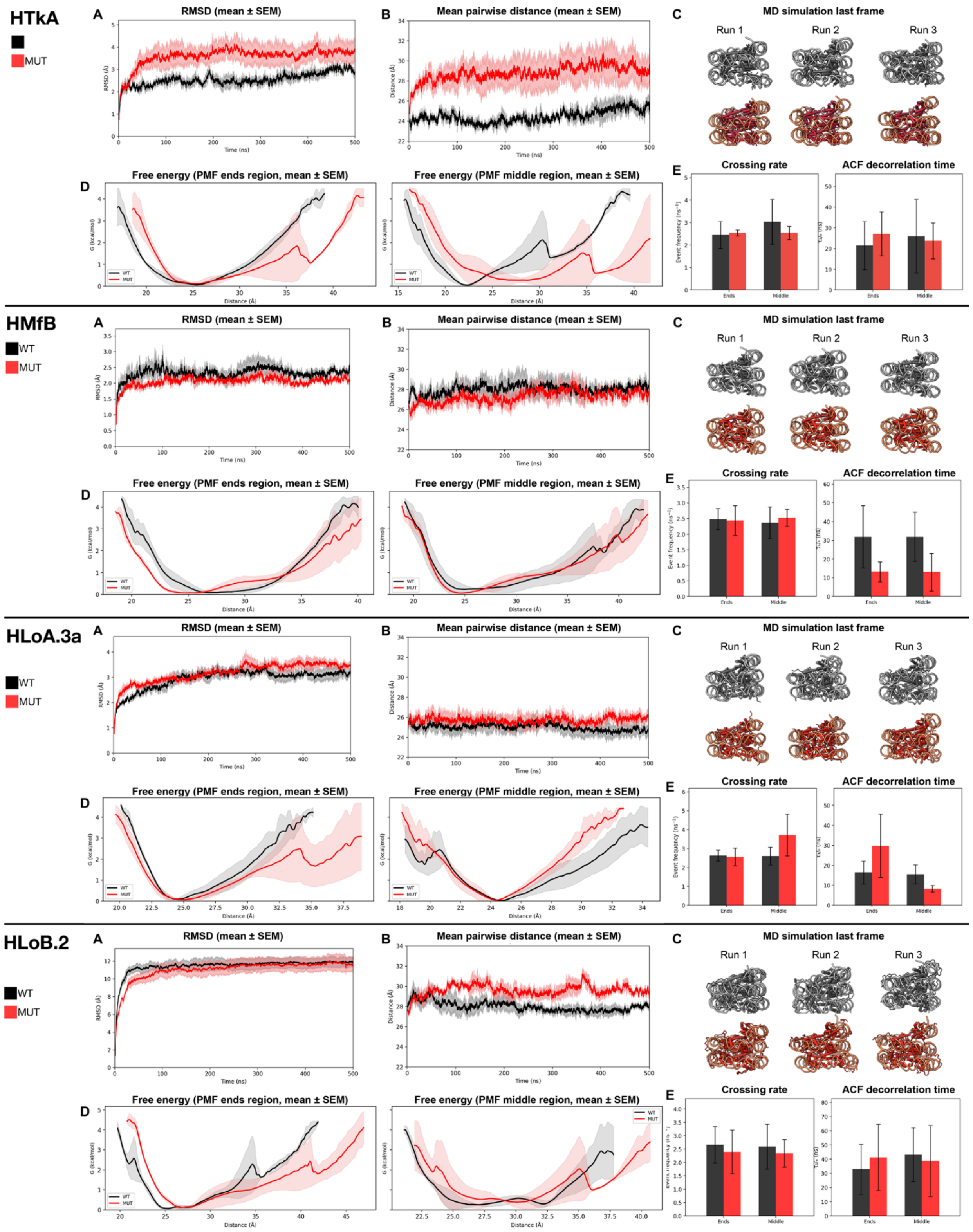
HTkA
MUT
A
RMSD (mean ± SEM)
B
Mean pairwise distance (mean ± SEM)
C
MD simulation last frame
Run 1
Run 2
Run 3
D
Free energy (PMF ends region, mean ± SEM)
Free energy (PMF middle region, mean ± SEM)
E
Crossing rate
ACF decorrelation time
HMfB
WT
MUT
A
RMSD (mean ± SEM)
B
Mean pairwise distance (mean ± SEM)
C
MD simulation last frame
Run 1
Run 2
Run 3
D
Free energy (PMF ends region, mean ± SEM)
Free energy (PMF middle region, mean ± SEM)
E
Crossing rate
ACF decorrelation time
HLoA.3a
WT
MUT
A
RMSD (mean ± SEM)
B
Mean pairwise distance (mean ± SEM)
C
MD simulation last frame
Run 1
Run 2
Run 3
D
Free energy (PMF ends region, mean ± SEM)
Free energy (PMF middle region, mean ± SEM)
E
Crossing rate
ACF decorrelation time
HLoB.2
WT
MUT
A
RMSD (mean ± SEM)
B
Mean pairwise distance (mean ± SEM)
C
MD simulation last frame
Run 1
Run 2
Run 3
D
Free energy (PMF ends region, mean ± SEM)
Free energy (PMF middle region, mean ± SEM)
E
Crossing rate
ACF decorrelation time

84

**Supplementary Figure 6. Effects of detected post-translation modifications (PTMs) on hypernucleosome structures in molecular dynamics simulations.** Histones HTkA from *T. kodakarensis*, HMfB from *M. fervidus*, HLoA.3a, and HLoB.2 from Loki-B35 were studied. All analyses compare complexes with wild-type protein (WT, black-grey) and mutants with all detected PTMs incorporated (MUT, red-pink). (a) RMSD curves. (b) Mean pairwise distances of 10 DNA pairs. (c) Snapshots of the last frame from three runs. (d) Free energy curves at the end and middle regions. (e) Crossing rate from the mean distance of WT and decorrelation time calculated from the autocorrelation function. All data is presented as a mean ± s.e.m. ($n$ = 3).

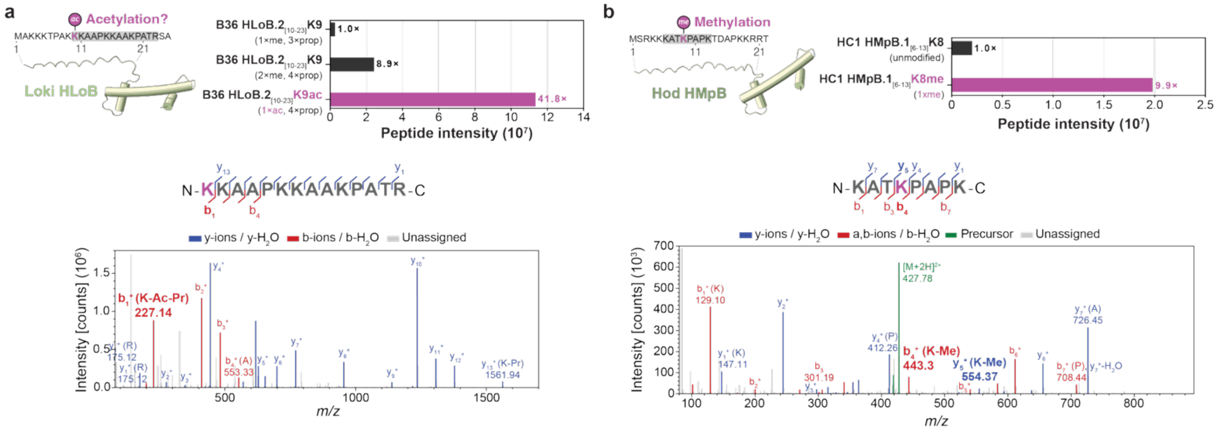


| # | a+ | a2+ | b+ | b2+ | Residue | y+ | y2+ | # |
|---|---|---|---|---|---|---|---|---|
| 1 | 199.14410 | 100.07569 | **227.13902** | 114.07315 | K-ac-pr | | | 14 |
| 2 | 383.26528 | 192.13628 | 411.26020 | 206.13374 | K-pr | 1561.94244 | 781.47486 | 13 |
| 3 | 454.30240 | 227.65484 | 482.29731 | 241.65229 | A | 1377.82126 | 689.41427 | 12 |
| 4 | 525.33951 | 263.17339 | 553.33443 | 277.17085 | A | 1306.78415 | 653.89571 | 11 |
| 5 | 622.39227 | 311.69978 | 650.38719 | 325.69723 | P | 1235.74703 | 618.37716 | 10 |
| 6 | 806.51345 | 403.76036 | 834.50837 | 417.75782 | K-pr | 1138.69427 | 569.85077 | 9 |
| 7 | 990.63463 | 495.82095 | 1018.62954 | 509.81841 | K-pr | 954.57309 | 477.79018 | 8 |
| 8 | 1061.67174 | 531.33951 | 1089.66666 | 545.33697 | A | 770.45191 | 385.72960 | 7 |
| 9 | 1132.70886 | 566.85807 | 1160.70377 | 580.85552 | A | 699.41480 | 350.21104 | 6 |
| 10 | 1316.83004 | 658.91866 | 1344.82495 | 672.91611 | K-pr | 628.37769 | 314.69248 | 5 |
| 11 | 1413.88280 | 707.44504 | 1441.87771 | 721.44250 | P | 444.25651 | 222.63189 | 4 |
| 12 | 1484.91991 | 742.96359 | 1512.91483 | 756.96105 | A | 347.20374 | 174.10551 | 3 |
| 13 | 1585.96759 | 793.48743 | 1613.96251 | 807.48489 | T | 276.16663 | 138.58695 | 2 |
| 14 | | | | | R | 175.11895 | 88.06311 | 1 |

| # | a+ | a2+ | b+ | b2+ | Residue | y+ | y2+ | # |
|---|---|---|---|---|---|---|---|---|
| 1 | 101.10732 | 51.05730 | 129.10224 | 65.05476 | K | | | 8 |
| 2 | 172.14444 | 86.57586 | 200.13935 | 100.57331 | A | 726.45085 | 363.72906 | 7 |
| 3 | 273.19212 | 137.09970 | 301.18703 | 151.09715 | T | 655.41374 | 328.21051 | 6 |
| 4 | 415.30273 | 208.15500 | **443.29764** | 222.15246 | K-me | **554.36606** | 277.68667 | 5 |
| 5 | 512.35549 | 256.68139 | 540.35041 | 270.67884 | P | 412.25545 | 206.63136 | 4 |
| 6 | 583.39261 | 292.19994 | 611.38752 | 306.19740 | A | 315.20268 | 158.10498 | 3 |
| 7 | 680.44537 | 340.72632 | 708.44029 | 354.72378 | P | 244.16557 | 122.58642 | 2 |
| 8 | | | | | K | 147.11280 | 74.06004 | 1 |

**Supplementary Figure 7. Additional MS/MS validation of histone tail post-translational modifications in Asgard archaea. a**, MS/MS spectrum and fragment ion table for the Loki-B36 HLoB tailed histone peptide KKAAPKKAAKPATR showing possible acetylation at K1 (HLoB$_{[10-23]}$K9ac). Propionylation (pr) of unmodified lysine residues (K2, K6, K7, K10) results from chemical derivatization during sample preparation. **b**, MS/MS spectrum and fragment ion table for the Hod-HC1 HMpB tailed histone peptide KATKPAPK showing methylation at K4 (HMpB.1$_{[6-13]}$K8me1). The diagnostic $b_4^+$ and $y_5^+$ ion pair localizes the modification to K4. In both panels, fragment ion abundance is plotted at various mass-to-charge ratios (*m/z*). The b and y ions and their losses of $H_2O$ are marked in red and blue, respectively; precursor ions in green; unassigned peaks in light grey. Nonessential ion labels have been omitted for readability. Fragment ion tables list theoretical *m/z* values for all detected ion series. ac, acetylation; pr, propionylation; me, methylation. Histone residues are numbered canonically from the second residue, as the initiator methionine is absent in the mature polypeptide.

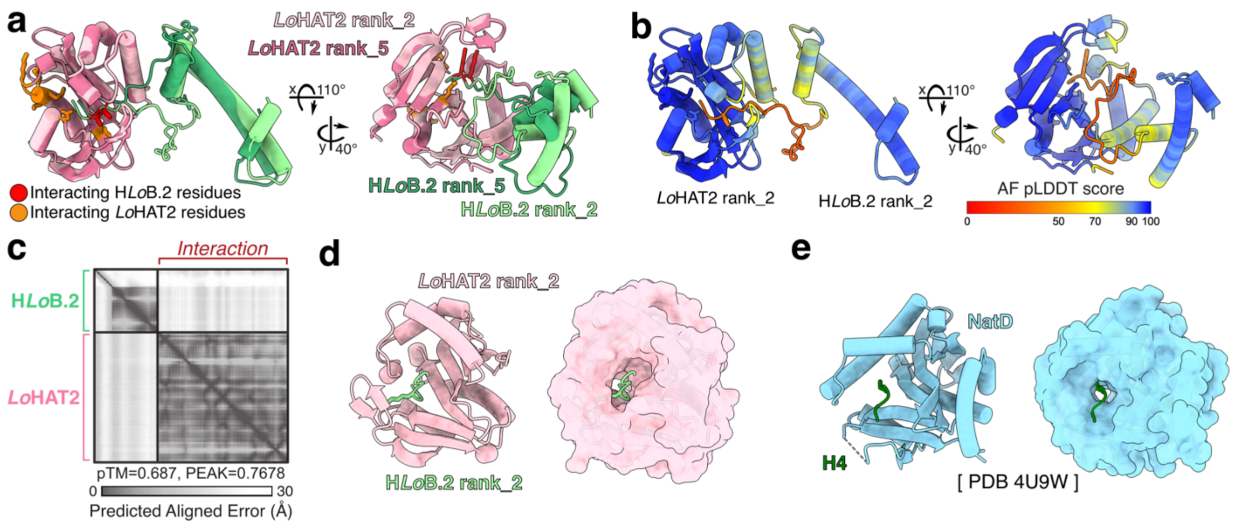


**Supplementary Figure 8. AlphaFold structural prediction of Asgard histone writer. a-d**, AlphaFold2 (AF2) predictions of *Lo*HAT2–HLoB.2 complexes. **a**, Superimposed AF2 models of two most distinct predictions (rank_2 and rank_5) of *Lo*HAT2 (two shades of pink) engaging the N-terminal tail of HLoB.2 (two shades of green) displays high prediction confidence in the protein core fold of both LoHAT2 and HLoB.2 together with their interaction interface. Relative position of histone is variable, due to highly flexible histone tail. As displayed also in Figure 3c, conserved catalytic interface residues on *Lo*HAT2 (orange: Gly106–Ser111 (GKGLMS), Glu95, Asn138) related to acetyl-CoA binding sites are positioned to contact the first accessible lysine on the histone tail (red: Lys3). Models shown in two orientations. **b**, Highest rank model from a, (rank_2) colored by the Predicted Local Distance Difference Test (pLDDT) score from AF2 and shown in two orientations. **c**, Predicted Aligned Error (PAE) plots from AF2 showing the *Lo*HAT2–HLoB.2 interaction prediction. **d**, Highest rank model from a, (rank_2) of *Lo*HAT2 and truncated HLoB.2 model showing the histone tail interacting residue (Lys3), displayed as atom, together with flanking residues. Cartoon representation (left) and surface representation of *Lo*HAT2 (right) visualizing the predicted catalytic cleft, where histone tail is binding. **e,** Experimentally determined structure of human N-terminal acetyltransferase type D (NatD) (light blue) and part of H4 tail (dark green) (PDB: 4U9W) showing the histone tail interacting residue (Ser1) displayed as atom. Cartoon representation (left) and surface representation of NatD visualize the predicted catalytic cleft, where the histone tail is binding, similarly as observed on the predicted *Lo*HAT2 model.

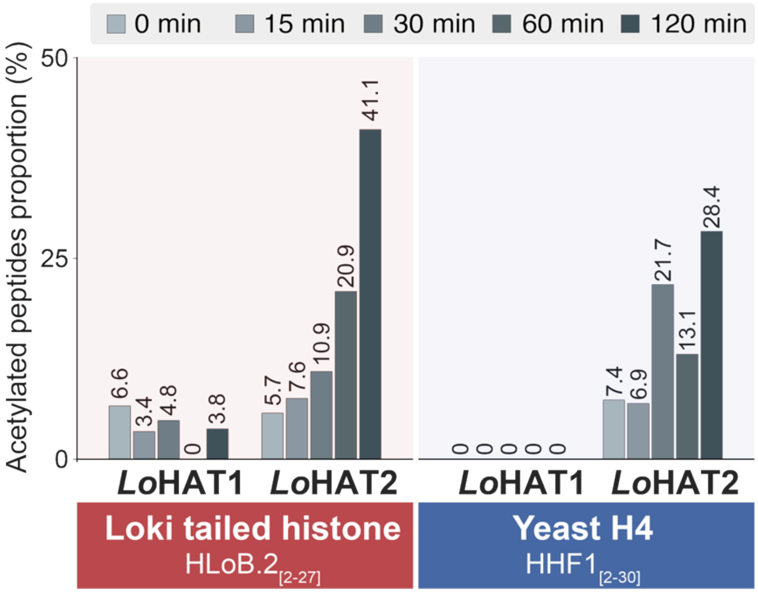


**Supplementary Figure 9. In vitro HAT activity assays using mass spectrometry**. In vitro HAT activity assays using synthetic peptides from the Loki histone tail (HLoB.$2_{[2-27]}$) or the yeast histone H4 tail (HHF$1_{[2-30]}$) as substrates were incubated with Loki HAT enzymes (*Lo*HAT1 and *Lo*HAT2). Mass spectrometry reveals that *Lo*HAT2 exhibits robust acetyltransferase activity, generating approximately a 10-fold higher acetylated peptide signal after 120 minutes compared to unincubated controls.

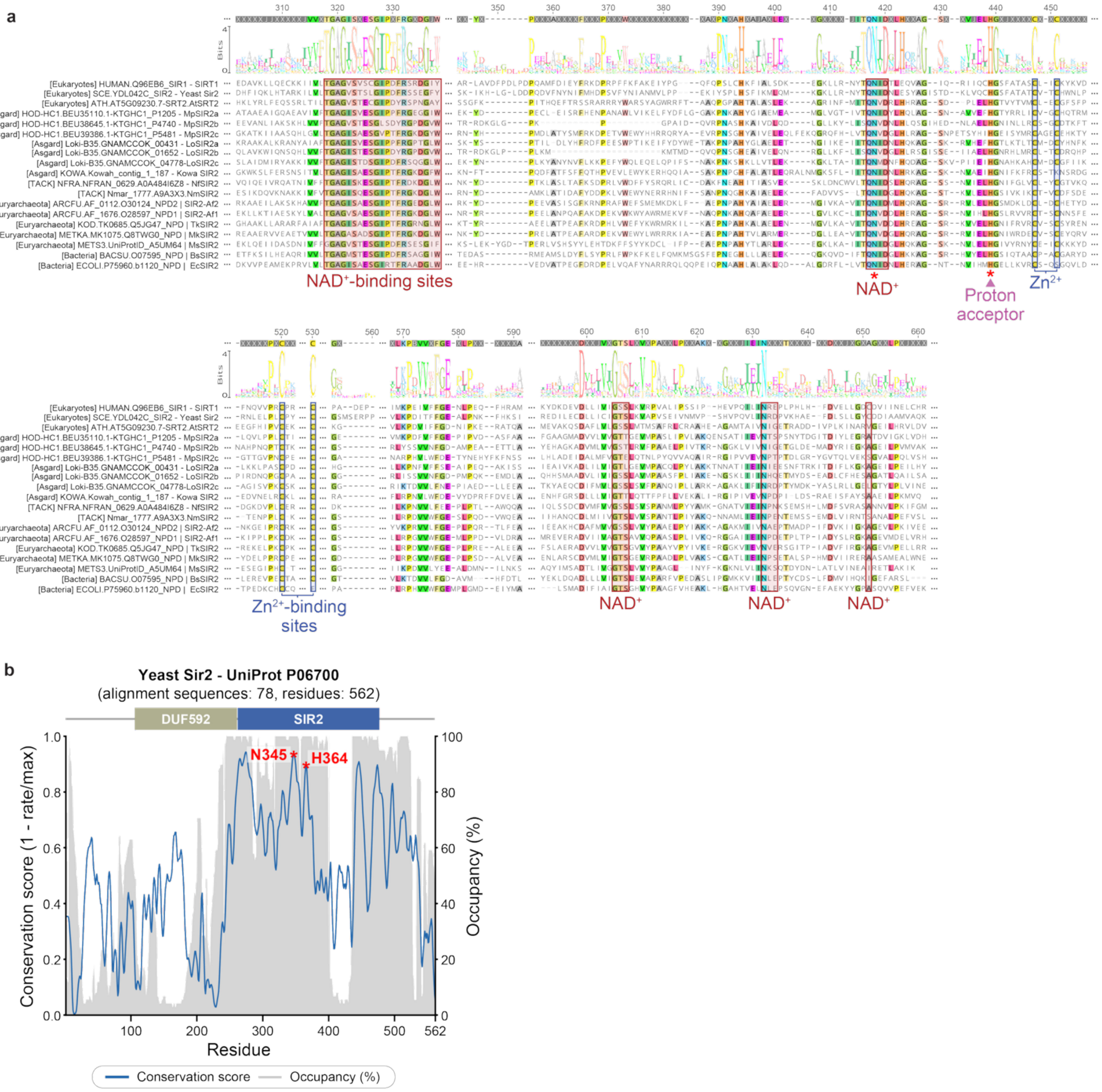


**Supplementary Figure 10. Multiple sequence alignment and conservation of SIR2 family deacetylases.** Top: Multiple sequence alignment of representative SIR2/sirtuin family NAD$^+$-dependent deacetylases from eukaryotes, Asgard archaea, TACK, Euryarchaeota, and Bacteria ($n$ = 19). Two alignment blocks are shown, spanning the catalytic core. NAD$^+$-binding residues (red boxes), proton acceptor residue (purple triangle), and Zn$^{2+}$-coordinating cysteines (blue boxes) are annotated. Red asterisks mark residues mutated to generate the catalytically inactive variant (*Lo*SIR2a-N106A/H123A). Consensus sequence logos above each block indicate positional conservation. Bottom panel: Residue-level conservation index mapped onto yeast Sir2 (UniProt: P06700). The blue line shows the conservation score (inverse Shannon entropy) across the 562 amino acid sequence. The grey line indicates alignment occupancy. Domain architecture is shown above (targeting domain DUF592, olive; SIR2 main domain, blue). Key catalytic residues N345 and H364 are labeled (red asterisks). The high conservation of catalytic and cofactor-binding residues across all domains of life supports an ancestral $NAD^+$-dependent deacetylase activity.

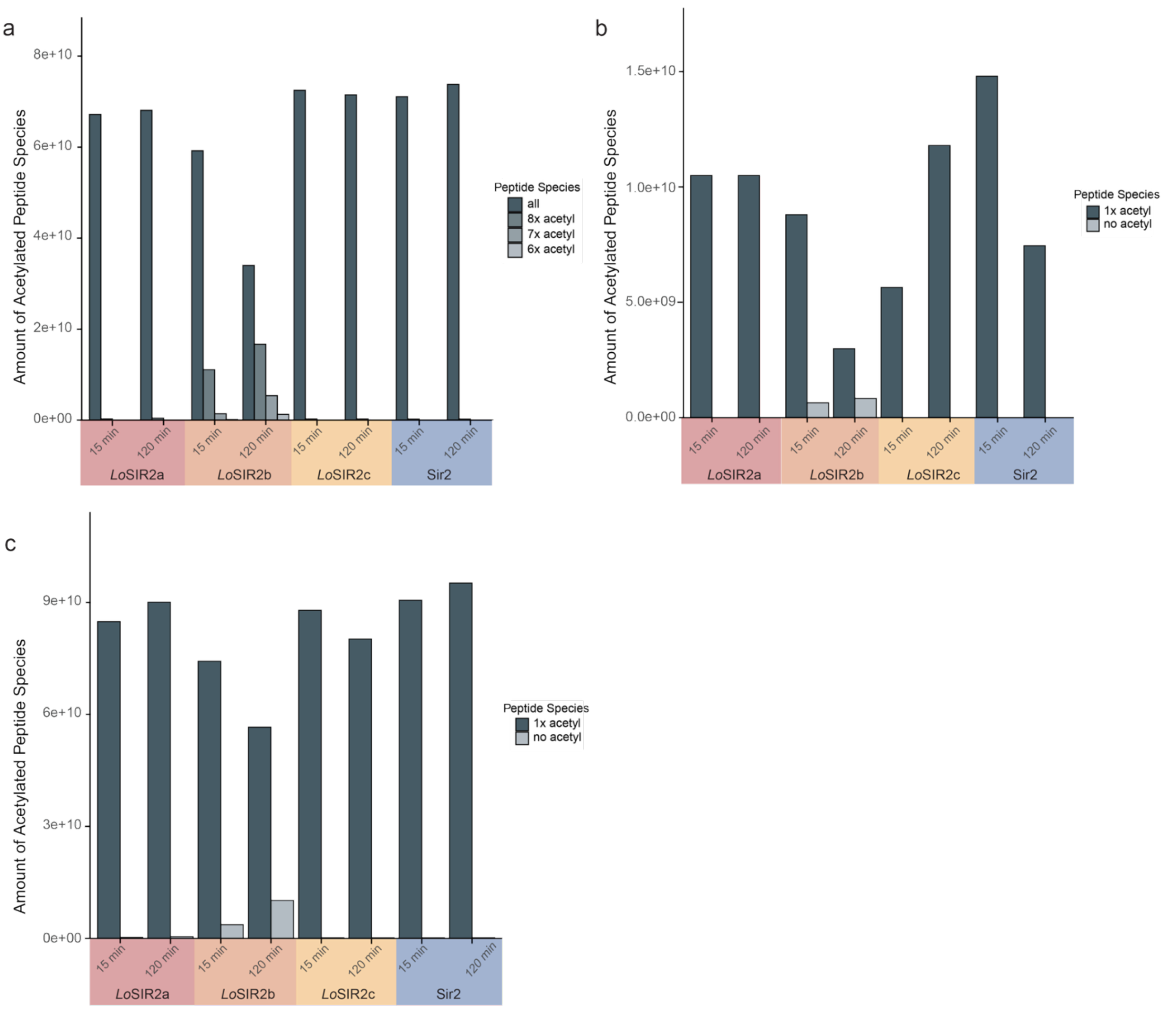


**Supplementary Fig. 11. In vitro SIR2 deacetylation assays quantified by mass spectrometry.** **a,** Deacetylation of fully acetylated Loki histone tail peptide HLoB.$2_{[2–27]}$ (all Kac). The abundance of acetylated peptides (all (9×), 8×, 7×, 6× acetyl) is shown for each enzyme at 15 and 120 min. *Lo*SIR2b progressively converts the fully acetylated substrate into lower acetylation states, whereas *Lo*SIR2a,c and yeast Sir2 show minimal activity on this substrate. **b,** Deacetylation of mono-acetylated Loki histone tail peptide HLoB.$2_{[2–27]}$K9ac. The abundance of mono-acetylated (1× acetyl) and unmodified (no acetyl) peptide is shown. *Lo*SIR2b efficiently removes the single acetylation with the unmodified peptides accumulating by 120 min. **c,** Deacetylation of yeast histone HHF1 tail peptide $H4_{[2–30]}$K16ac. *Lo*SIR2b deacetylates the yeast substrate, whereas other SIR2s show little or no detectable activity under these conditions. In all panels, peptide abundance (y-axis) was measured by mass spectrometry. Coloured backgrounds group enzymes by origin: *Lo*SIR2 paralogs (red, orange, yellow) and yeast Sir2 (blue).

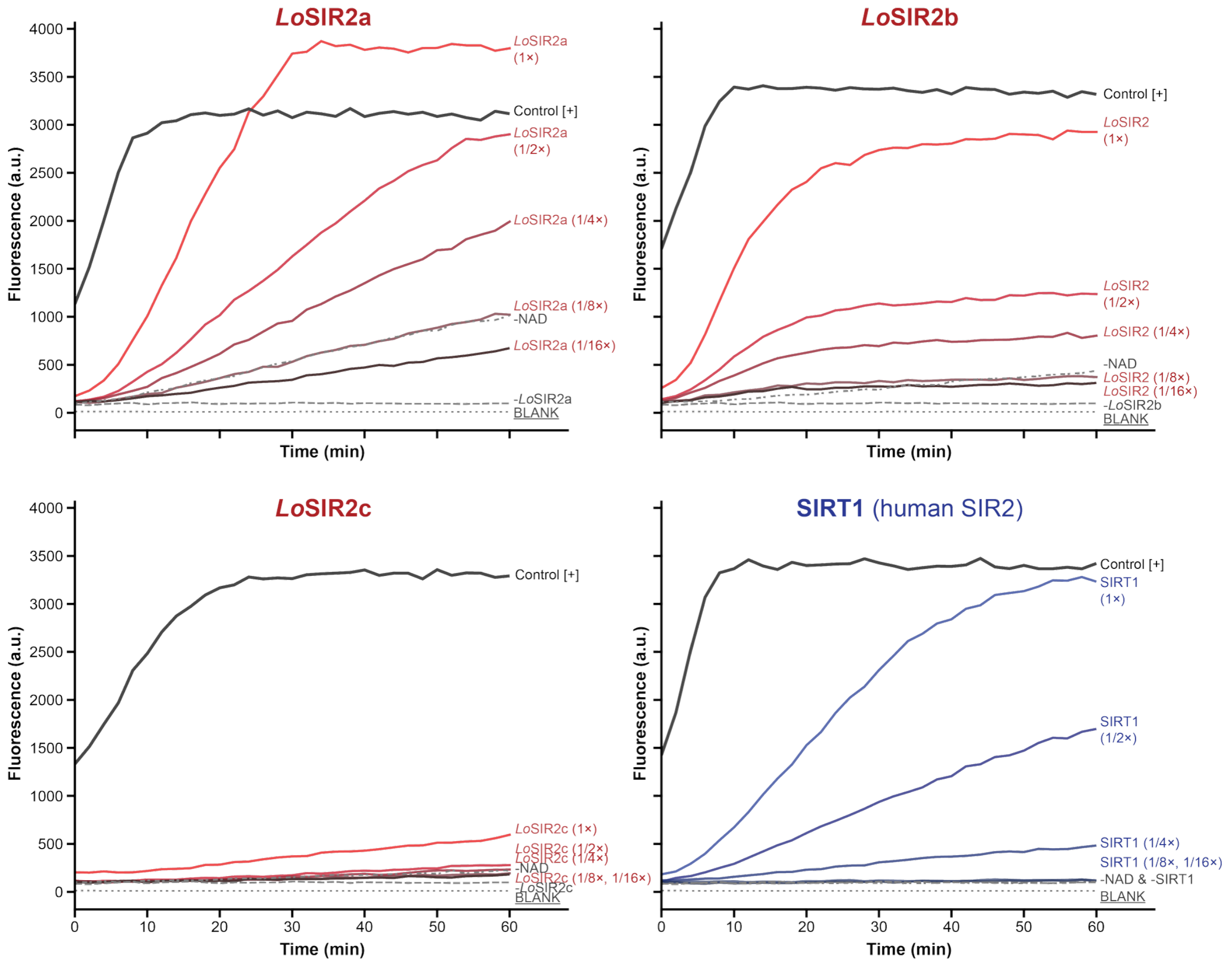


**Supplementary Figure 12. Enzymatic activity of recombinant Asgard archaeal SIR2 deacetylases.** Fluorescence-based deacetylase activity assays were performed for recombinant SIR2 enzymes from Asgard archaea (*Lo*SIR2a, *Lo*SIR2b, *Lo*SIR2c) and human SIRT1 (positive control). Fluorescence (arbitrary units, a.u.) is plotted over time (0–60 min) for varying enzyme concentrations (1× = 4 µg; 1/2×, 1/4×, 1/8×, 1/16×). The 1× concentration corresponds to ~1.4 µM for *Lo*SIR2a–c (His6-GST fusions, MW ~56 kDa) and ~1.0 µM for human SIRT1 (UniProt Q96EB6, MW ~83 kDa). All measurements were performed in duplicates, and data are presented as mean values. *Lo*SIR2a and *Lo*SIR2b display robust, dose-dependent deacetylase activity comparable to human SIRT1, with fluorescence increasing over time and plateauing at higher enzyme concentrations. *Lo*SIR2c shows minimal activity under standard assay conditions but produces a signal in the deacetylated peptide positive control ('#3', dashed grey line), suggesting that the lack of activity may be due to problems with recombinant protein folding or stability rather than an inherent absence of catalytic function. Negative controls (no $NAD^+$ and no enzyme blank) show no fluorescence increase, confirming that the observed activity is $NAD^+$-dependent.
